# Multiple groups of neurons in the superior colliculus convert value signals into saccadic vigor

**DOI:** 10.1101/2025.06.24.661386

**Authors:** Atul Gopal, Okihide Hikosaka

## Abstract

Eye movements directed to high-valued objects in the environment are executed with greater vigor. Superior Colliculus (SC) - a subcortical structure that controls eye movements - contains multiple subtypes of neurons that have distinct functional roles in generating saccades. How does value-related information processed in other parts of the brain affect the responses of these different subtypes of SC neurons to facilitate faster saccades? To test this, we recorded four subtypes of neurons simultaneously while the monkey made saccades to objects they had been extensively trained to associate with large or small rewards (i.e., good or bad). In three subtypes of neurons (visual, visuomotor, and motor), the good objects elicited more spikes than bad objects. More importantly, using a bootstrapping procedure, we identified three separable phases of activity: 1) early visual response (E_VIS_), 2) late visual response (L_VIS_), and 3) pre-saccadic (Pre_SAC_) motor response in these neuronal subtypes. In each subtype of neurons, the value of objects (good vs. bad) was positively correlated with the activity in the L_VIS_ and Pre_SAC_ phases but not the E_VIS_ phase. These data suggest that the value information from other brain regions modulates the visual (L_VIS_) and the motor (Pre_SAC_) responses of visual, visuomotor, and motor neurons. This enhanced activation facilitates the faster initiation and execution of the saccade based on the value of each object. In addition, we found a novel class of tonically active neurons that decrease their activity in response to object onset and remain inhibited till the end of the saccade. We suggest that these tonic neurons facilitate the saccade to objects by disinhibiting the interactions between the other three SC neurons.

## INTRODUCTION

Reward is a critical factor that shapes the behavior of all organisms. Positive outcomes encourage animals to approach or explore objects and lead to invigorated action, whereas negative consequences lead to avoidance and slower actions (Dickinson & Balleine, 1994; Watson & Platt, 2008). The influence of reward can be seen even in simple saccadic eye movements, such that reaction times are faster and peak velocities are higher for saccades to high-value objects and/or locations (Reppert et al., 2015; Takikawa et al., 2002; Yoon et al., 2020). This behavioral modulation by reward necessitates an interaction between value processing networks and motor control networks within the brain, which enable the integration of motivational information with movement planning and execution.

Saccadic eye movements are planned and executed by a network of brain regions, including the frontal eye fields, parietal cortex, and superior colliculus (SC) (Munoz, 2002; Pouget, 2014). In this pathway, SC is the final critical node that can serve as an interface between cognitive signals and movement planning (Basso et al., 2021; Basso & May, 2017) prior to the generation of the motor commands by the brainstem circuits to initiate saccades (Munoz et al., 2000; David L. Sparks, 2002). The SC receives sensory input and transforms it into motor commands that drive eye movements with the help of three distinguishable neuronal cell types-visual, visuomotor, and motor neurons (Wurtz & Albano, 1980; Wurtz & Optican, 1994). These neurons are differentially localized in the superficial and intermediate layers of SC, forming a local circuit. The activity of these neurons controls saccade initiation and execution which can be measured as reaction time and peak velocity, respectively (Dorris et al., 1997; Katnani & Gandhi, 2012; R. J. Krauzlis, 2003; C. Lee et al., 1988; Waitzman et al., 1991).

The brain contains a large network of cortical and subcortical regions to process and assign value from the incoming visual inputs. The basal ganglia is a critical node in this network and is particularly well-documented for its role in influencing saccadic eye movements (Hikosaka et al., 2006). The neural activities in multiple basal ganglia (BG) nuclei, including the caudate nucleus (Kawagoe et al., 1998; H. F. Kim & Hikosaka, 2013), globus pallidus (Hong & Hikosaka, 2008; H. F. Kim et al., 2017), and substantia nigra (Sato & Hikosaka, 2002; Yasuda & Hikosaka, 2015) are known to differentiate between high-valued (good) and low-valued (bad) objects/positions. This value information is then transmitted from the output nuclei of BG, the substantia nigra pars reticulata (SNr), to other brain regions dedicated to controlling saccadic behavior, such as the SC (Hikosaka et al., 2000). Previous studies have shown that BG input to SC takes the form of disinhibition (Hikosaka & Wurtz, 1983; Liu & Basso, 2008; Yasuda & Hikosaka, 2017) mediated by the SNr neurons. Additionally, SNr neurons are modulated by the value of objects present in the contralateral visual field, reducing its activity for good objects and enhancing it in the case of bad objects (Amita et al., 2020; Yasuda & Hikosaka, 2015). Thus, these SNr neurons can interface with the visuomotor circuits in the SC and gate the behavior by permitting saccades to valuable objects in the environment.

In addition to BG, many other cortical regions are also involved in processing value information and are able to provide value-related inputs to the neurons in SC. Previous studies have shown that neurons in the frontal eye fields are modulated by the expected value of the target prior to the action (X. Chen et al., 2020; Ding & Hikosaka, 2006; Glaser et al., 2016). Similarly, studies done in supplementary eye fields (Roesch & Olson, 2003; So & Stuphorn, 2010; Uchida et al., 2007) as well as in the lateral interparietal area (Leathers & Olson, 2012; Louie & Glimcher, 2010) have also shown that neurons in these regions are modulated by the expected reward at the end of the trial. Previous work from the lab using fMRI found regions across the brain that are modulated by value information (Ghazizadeh et al., 2018). More importantly, many of these cortical regions are known to have direct or indirect projections to SC (Borra et al., 2014; Helminski & Segraves, 2003; Lock et al., 2003; Paré & Wurtz, 2001) and may influence the saccadic behvaior. Value-sensitive inputs from these cortical and/or subcortical sources may interface with the local visuomotor circuits in SC formed by the multiple functional classes of neurons, affecting their responses. Understanding the dynamics of these interactions is crucial for a comprehensive mechanistic view of how reward shapes saccade behavior.

Previous studies have shown that SC neurons are modulated by reward expectation using an asymmetrically rewarded 1-DR task (Ikeda & Hikosaka, 2003, 2007) in which saccades to one direction were rewarded more than others in a block of trials. The enhanced activation was seen in some neurons when a target was placed in the location where the monkey was anticipating a high reward during the pre-target baseline and the initial transient phase of the post-target response. More recently, (Griggs et al., 2018) investigated the effect of value on the responses of visual neurons in the SC by using visual objects that were previously associated with higher or lower volumes of juice by repeated, consistent exposure for more than ten days. After the learning phase, the same objects were flashed in the receptive field (RF) of the SC neurons while the animal fixated at a central point. Visual neurons in the superficial layers of SC were activated during this passive peripheral viewing of the objects and responded with a higher firing rate for high-valued good objects than low-valued bad objects. This study used a passive fixation task to record only visual neurons of SC; hence, the impact of learned value on different neuronal subtypes that form the local circuitry in SC and their effect on saccade behavior was not fully assessed.

In this study, we compared the activity of different functional subtypes of SC neurons recorded using a linear array while the animals performed visually guided saccades towards high-valued good vs. low-valued bad objects. In contrast to single electrode recordings, a linear array allows the monitoring of neurons in different layers of SC simultaneously and thus provides a complete picture of the local SC circuit. The recorded neurons were broadly classified into four functional subtypes. High-valued good objects elicited more spikes than low-valued bad objects in three of the four subtypes of SC neurons, ∼ 100ms after target onsets. We also found that the stronger activation in these neurons leads to enhanced behavioral responses - lower reaction time (RT) and higher peak velocity (PV) - seen when saccades are directed to good objects.

## RESULTS

The SC contains multiple functional subtypes of neurons that play distinct roles in generating saccades. The value information from other cortical regions (e.g., FEF, LIP) and subcortical regions (e.g., SNr in the BG) is thought to interface with these neuronal subtypes and modulate their response properties to affect the saccadic responses to valuable objects. Here, we investigate how the response profiles of these different SC neurons are affected by reward, thereby modulating the RT and PV of the upcoming saccade. We tested this by recording SC neurons while the animals performed a visually guided saccade task (Fig-1A) towards fractal objects (Yasuda et al., 2012) previously associated with higher or lower volumes of juice reward (Fig-1B) presented in the contralateral visual field. A linear array with 24 contact points was positioned in the SC through a posteriorly implanted recording chamber, and multiple subtypes of neurons were simultaneously recorded from different layers (Fig-1C). We used the recorded activity in different time windows (Fig-1D) to classify the SC cells into four functional subtypes (See methods). In each recording session, the monkeys made saccades to fractal targets appearing at a specific eccentricity based on the identified RF of the recorded SC neurons (Fig-1E). Across sessions, targets appeared at eccentricities ranging from 4-25 degrees, and monkeys, on average, made 56 (min of 20 to max of 86) saccades to the good and 43 (min of 13 to max of 66) saccades to the bad objects.

**Figure 1:**
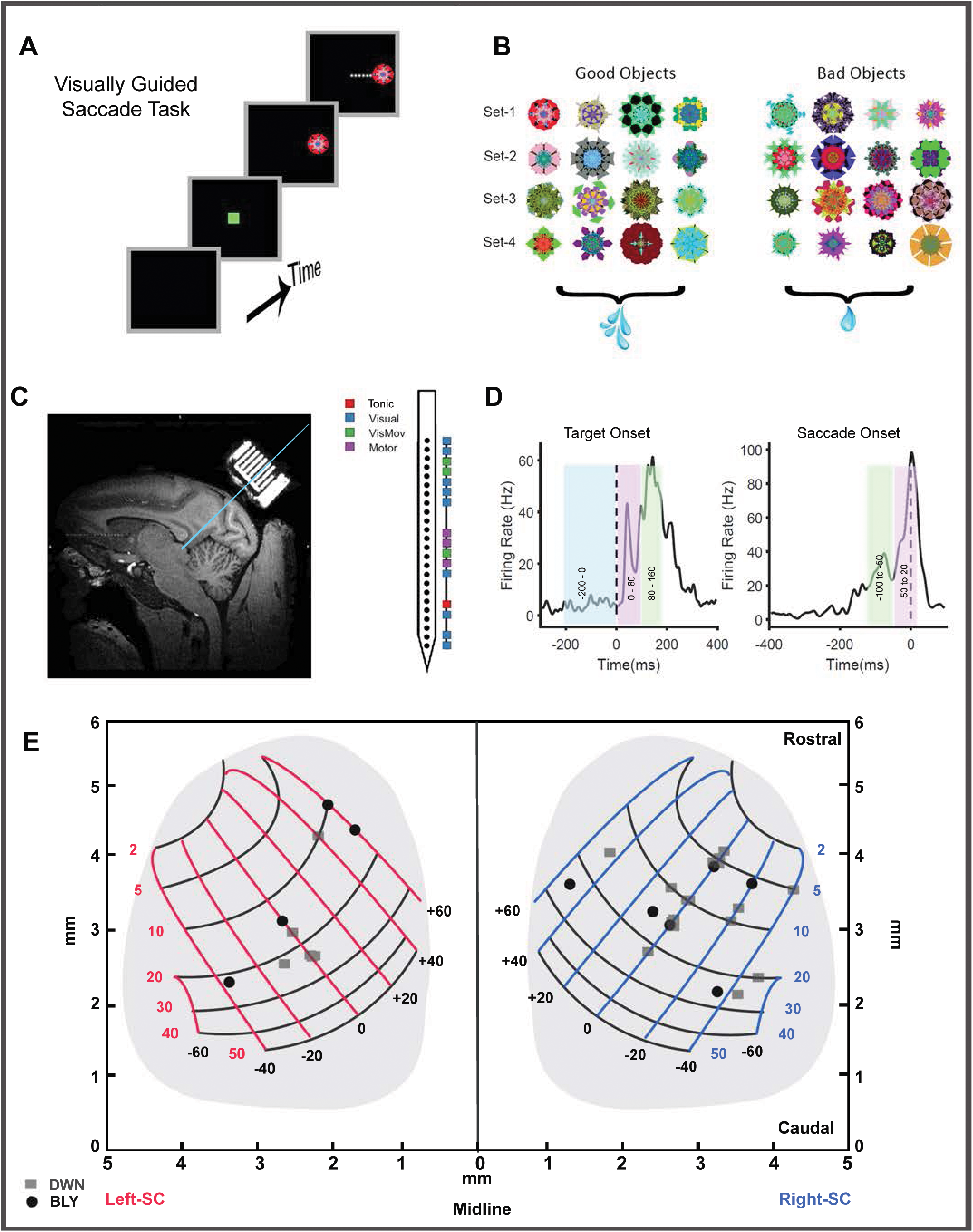
Methods used in this study. **(A)** Progression of events in a single trial of the visually guided saccade task. **(B)** The four object sets (eight fractals in each) were used as stimuli in the task. In each set, four objects were associated with a larger volume of juice (good objects), and four objects were associated with a lower volume (bad objects). Animals learn the value of good and bad objects through repeated exposure to the task. **(C)** A sagittal section of the brain from monkey DWN, visualized by MRI, showing the location of the posterior recording chamber and grid. A recording track targeting the right hemisphere of SC is shown as a blue line. The different subtypes of SC neurons recorded simultaneously using a linear array during an example session are also shown as a schematic. **(D)** The neural activity aligned on target onset (left) and saccade onset (right) is shown for a representative neuron. The firing rates in these time intervals (colored patches) were quantified and used to classify the SC cells into different subtypes. (E) A map showing the potential anatomical locations of recordings in SC derived based on the receptive field of the recorded neurons over 33 separate sessions. Sessions recorded from monkey DWN are shown in grey squares and from monkey BLY in black circles.

### Behavioral modulation of saccades

First, we tested if the saccadic eye movements in this task confirmed the previously reported reward-based behavioral modulation (Takikawa et al., 2002; Xu-Wilson et al., 2009). We separately computed the RT distribution of saccades to good and bad objects for each recording session (Fig-2A). In 26 out of 33 sessions, the RT distribution for good objects significantly differed (unpaired t-test, p < 0.045) from those for bad objects. The mean good object RT across all sessions (186 ± 25 ms) was significantly lower than the bad RT (211 ± 36 ms) (Fig-2D paired t-test, n=33, p<0.001), corroborating that saccade initiation is strongly modulated by reward.

**Figure 2:**
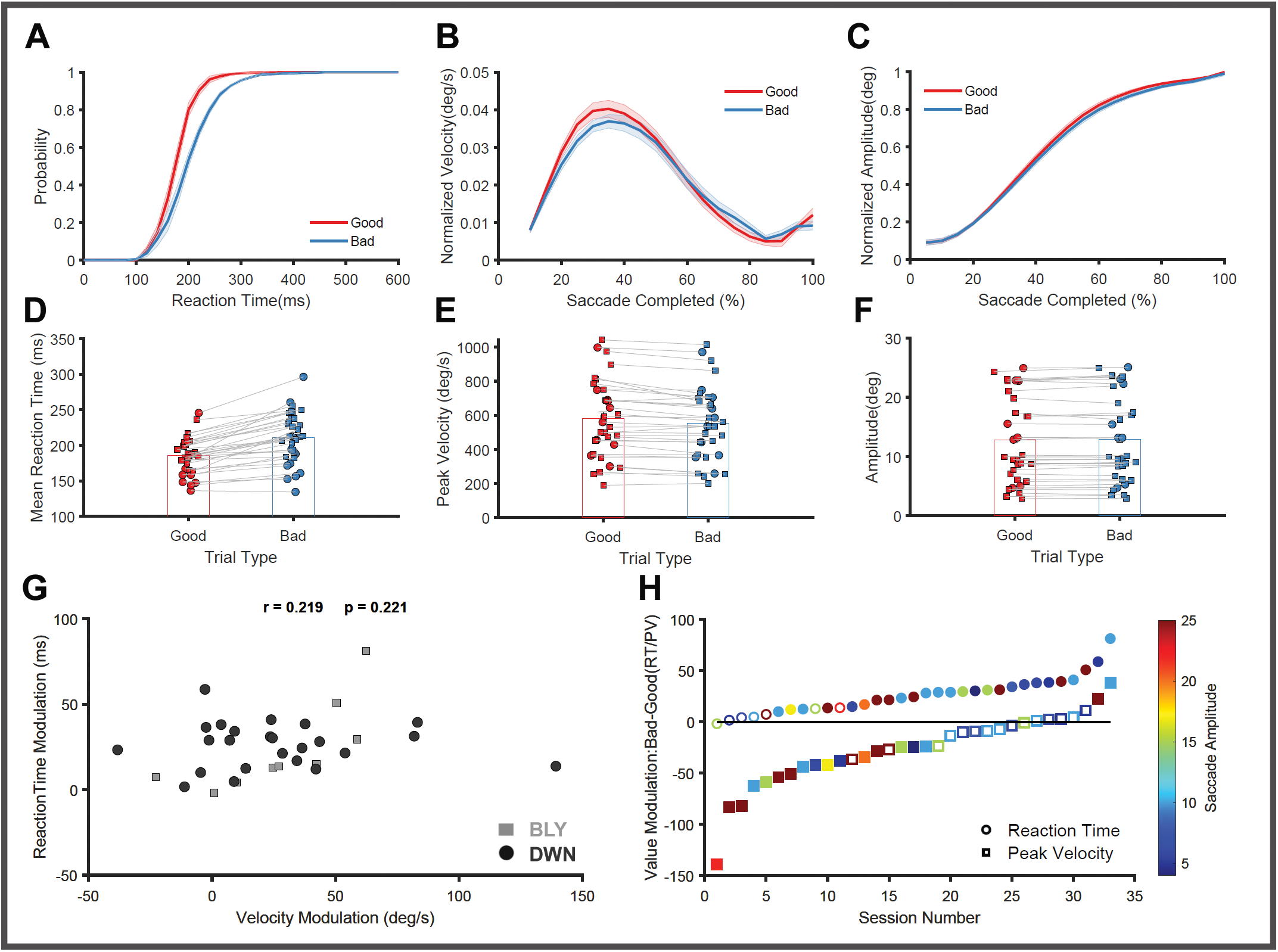
Effect of reward on saccadic behavior Saccadic behavior towards high-valued good (red) and low-valued bad (blue) objects is shown. **(A)** Average cumulative reaction time distributions of good and bad object trials from all the recording sessions are shown with the standard error of means as the colored envelope. **(B)** Average normalized velocity profiles of saccades directed to good and bad objects are shown with the standard error of means as the colored envelope. **(C)** Average normalized saccadic displacement profiles for good and bad trials are plotted separately. The mean RT **(D)**, mean PV **(E)** and mean amplitude **(F)** calculated separately for good and bad objects in each session (connected by the light grey line) are shown to depict the variability in behvaior across different recording sessions. The data obtained from the two animals are shown separately using different symbols. Bar graphs denote the population mean calculated by averaging across all the 33 recorded sessions. **(G)** A scatterplot between the value modulation (difference between good and bad object trials) observed in RT and PV across all the recorded sessions. The data obtained from the two animals are shown separately using different symbols. **(H)**. The extent of value modulation seen in RT (circle) and PV (square) across all the recorded sessions is plotted after sorting them in increasing order. The range of saccade amplitudes the animals made in different sessions of this experiment is shown as a colormap ranging from cool to warm colors. The sessions showing significant value modulation based on an unpaired t-test are shown as filled symbols.

Next, we calculated the average velocity profiles in each session separately for saccades to good and bad objects to test the effect of reward on saccade execution. The peak velocity of saccades towards good objects was significantly greater than saccades to bad objects in 18 out of 33 sessions (unpaired t-test, p<0.04). The mean good object PV (740 ± 321 deg/sec) was significantly greater than the mean bad object PV (700 ± 300 deg/sec) (Fig-2E paired t-test, n=33, p<0.001). During the execution of saccades, peak velocity is affected by the intended amplitude through the main sequence relationship (Bahill et al., 1975; Bollen et al., 1993). Here, monkeys made saccades to different amplitudes ranging from 4 to 25 degrees, and there is a strong correlation between the PV and the amplitude (Pearson’s correlation =0.83 p < 0.001). Hence, we normalized the velocity profiles as described in the methods and plotted them separately (Fig-2B) to show that saccades to good objects were executed faster compared to bad objects. In addition, we also tested if the difference in PV observed during saccades to good and bad objects is due to changes in saccade amplitudes. There is no significant difference in the average saccade displacement profiles (Fig-2C KS Test, p>0.05) in all of the 33 recorded sessions or in the average amplitude (Fig-2F paired t-test, n=33, p=0.16) observed between the good (12.78 ± 7.2 degree) and bad (12.89 ± 7.36 degrees) object saccades; hence the PV modulation observed in saccades is independent of the amplitude. This further corroborates that in this dataset, the reward associated with good and bad objects can modulate the execution of saccadic eye movements, as shown previously.

We further tested if the modulation observed in RT and PV are related to each other by plotting the difference in RT between good and bad objects against the difference observed in PV across the different sessions. There is no clear evidence of a systematic relationship (Fig-2G Pearson’s correlation = 0.2; p=0.22) between RT and PV modulations, suggesting that reward may independently affect these behavioral metrics. Further, we also checked if the different target eccentricities used in this study have any systematic effect on the observed reward modulation of saccade behavior (Fig-2H). Interestingly, sessions with larger target eccentricities showed stronger modulation of saccadic peak velocity (Pearson’s correlation = 0.47; p=0.005), while there was no effect of eccentricity on the observed RT modulation (Pearson’s correlation =-0.12; p=0.51).

### Reward Modulation in SC neurons

Next, we analyzed the responses of the SC neurons while the animal directed their saccades to the good and bad objects (Fig-S1 for representative neurons, Fig-3 for population activity). We analyzed 520 neurons recorded from two monkeys, which were then divided into four functional subtypes (See methods for classification). The population responses were calculated separately for each functional subtype by averaging across all the recorded visual (n=239), visuomotor (n=89), motor (n=96), and tonic neurons (n=89) aligned on target onset as well as saccade onset (Fig-3).

**Figure 3:**
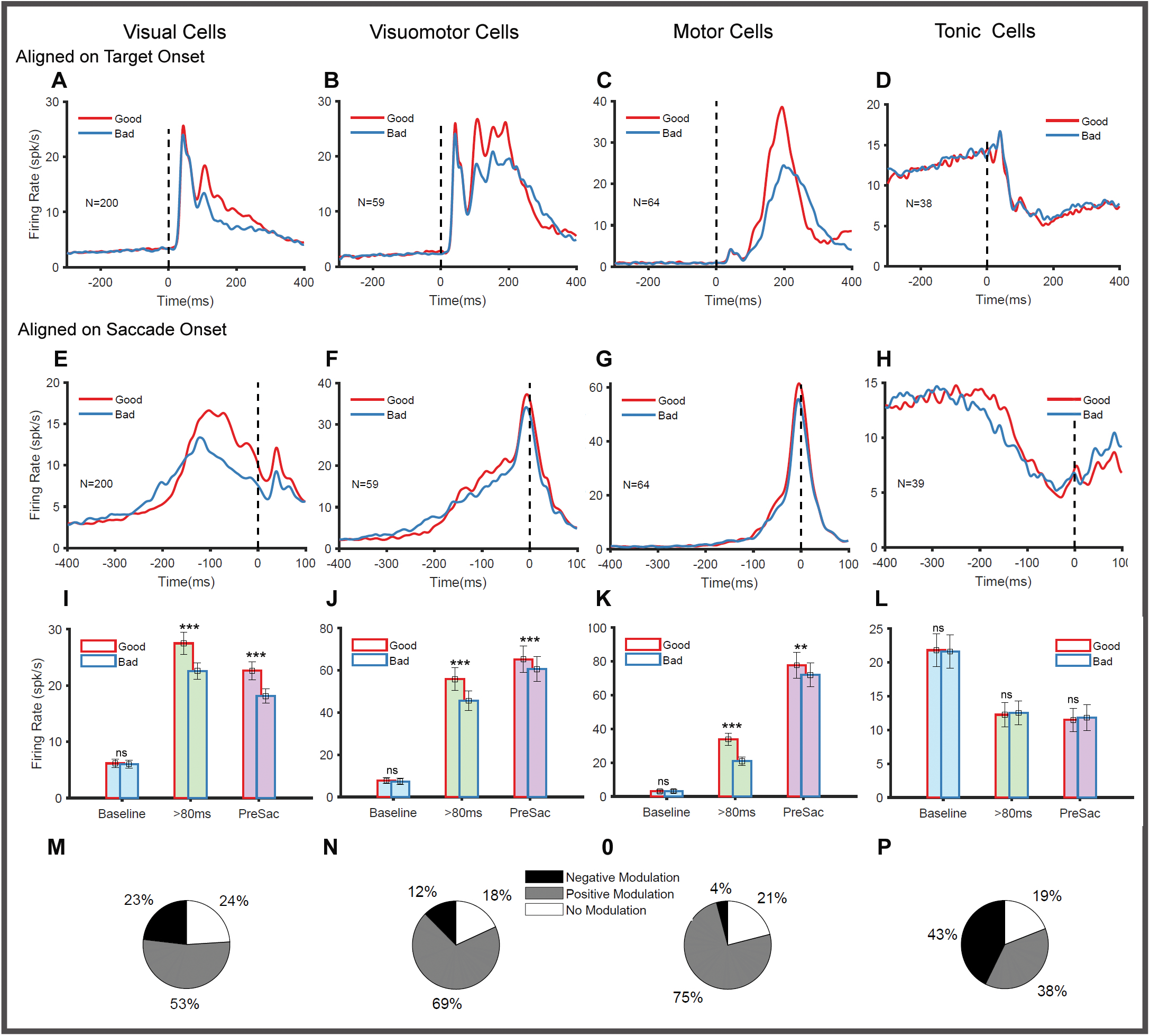
Population response of SC neurons to reward modulation The average SC population response aligned on stimulus onset when saccades were directed to the good objects (red) and bad objects (blue) in the RF is shown separately for visual **(A)**, visuomotor **(B)**, motor **(C)**, and tonic **(D)** neurons. The population response of the SC neurons, when aligned on saccade onsets, is shown separately for visual **(E)**, visuomotor **(F)**, motor **(G)**, and tonic **(H)** neurons. The firing rate calculated in the time intervals-300 to 0 (light blue) before the target onset, greater than 80ms (light green) aligned on the target onset, and-50 to 20ms aligned on the saccade onset (light violet) are shown as bar graphs separately for the good (red outline) and bad (blue outline) trials. The error bars denote the standard error of means. The firing rate is separately quantified for the visual **(I)**, visuomotor **(J)**, motor **(K)**, and tonic **(L)** neurons. The piecharts below show the proportion of positively modulated (grey), negatively modulated (black), and non-modulated (white) neurons within the populations of visual **(M)**, visuomotor **(N)**, motor **(O)**, and tonic **(P)** neurons. *** denotes p<0.001 ** denotes p<0.005 to p>0.001 and ns denote p>0.05.

Visual neurons exhibit an initial transient increase in their firing with a latency of 20-40 ms when an object appears in their RF (Fig-3A). At about 80 ms after target onset, their responses, which are sustained above the baseline, became larger to good objects (27 spks/s) than bad (22 spks/s) objects (paired t-test p<0.001). When aligned on saccade onset (Fig-3E), we did not observe any pre-saccadic burst in these neurons; instead, the activity was maintained stably above the baseline with higher (paired t-test, p<0.001) activation for good objects (22 spks/s) compared to bad objects (18spks/s). This pattern of higher activation for good objects (positive modulation) is seen in 53% (n=133) of the recorded neurons. Interestingly, a minority of visual neurons, 23% (n=54), showed the opposite negative modulation, i.e., higher activation for bad objects. There were also 24% (n=56) of visual neurons that showed no significant difference in the activity between good and bad object saccades (Fig-3M).

Visuomotor neurons (Fig-3B) showed a clear excitatory phasic response to target onset, lasting until 80ms. These neurons maintained a higher level of activity for good objects (56 spks/s) in the time bins greater than 80 ms compared to bad objects (46spks/s). A clear excitatory presaccadic burst (Fig-3F) is seen in these neurons when the activity is aligned with saccade onset and this burst was positively modulated for good objects (FR_Good_ = 65spks/s compared to FR_Bad_ = 60 spks/s paired t-test, p = 0.003). Most visuomotor neurons were positively modulated 70% (n=61), while the negatively modulated cells constituted only 12% (n=11). The remaining 18% (n=16) of the visuomotor neurons showed no significant difference between the good and bad object conditions (Fig-3N).

Motor neurons, on the other hand, had a weak visual response (Fig-3C) but a clear excitatory burst prior to the motor onset (Fig-3G). These neurons responded with higher magnitude (Fig-3K paired t-test p < 0.001) when the ensuing saccade was directed to good objects (29 spks/s) compared to bad objects (19 spks/s). Their value-coding was less pronounced and restricted to the pre-saccadic period (Fig-3G) when aligned on saccade onset (FR_Good_ = 72.0spks/s compared to FR_Bad_ = 67.4 spks/s paired t-test with p=0.05). Similar to visuomotor neurons, most (75% n= 71) of the motor neurons were also positively modulated, and only a small minority 4% (n=4) were negatively modulated. The remaining 21% of the motor neurons (n=20) showed no difference in activity between good and bad object conditions (Fig-3O).

### Tonic Neurons: A new class of SC neurons

Unlike the other three groups of SC neurons, tonic neurons had significant activity (22 spks/sec) during the inter-trial interval and fixation duration. (Fig-4A). In response to an object (good or bad) presented in their RF, their activity was reduced to (12 spks/sec) at a latency of ∼40-60 ms. (Fig-4A,C (left) unpaired t-test, p<0.001). This inhibition persisted at least until the saccade onset and started to increase post-saccade (Fig-4B). To the best of our knowledge, these types of tonic SC neurons have not been reported previously. Earlier studies have identified a group of neurons in the rostral part of SC with foveal receptive fields and reduced their activity ∼40ms prior to the initiation of saccades (Munoz & Wurtz, 1992, 1993). This pause in their activity is required to remove the fixation and initiate saccadic eye movements. The tonic neurons identified here have markedly different properties from those of these foveal neurons in the rostral SC (R. J. Krauzlis, 2003; R. J. Krauzlis et al., 2000). Unlike rostral foveal neurons, the tonic neurons have eccentric receptive fields ranging from 4 to 25 degrees (Fig-4D). They were not specifically localized in the rostral zone of the SC and were found in recording tracts in the caudal part of the SC (Fig-1E). More importantly, unlike foveal neurons, which are modulated by the onset/offset of the central fixation cue, most tonic neurons (77/88) are unaffected by the onset of the fixation cue. There was no significant difference in the activity recorded before (15 spks/s) and after (16 spks/s) the onset of the fixation cue (Fig-4C (right), unpaired t-test p=0.754). Given these, we suggest that the tonic neurons are a separate class of SC neurons that have not previously been reported.

**Figure 4:**
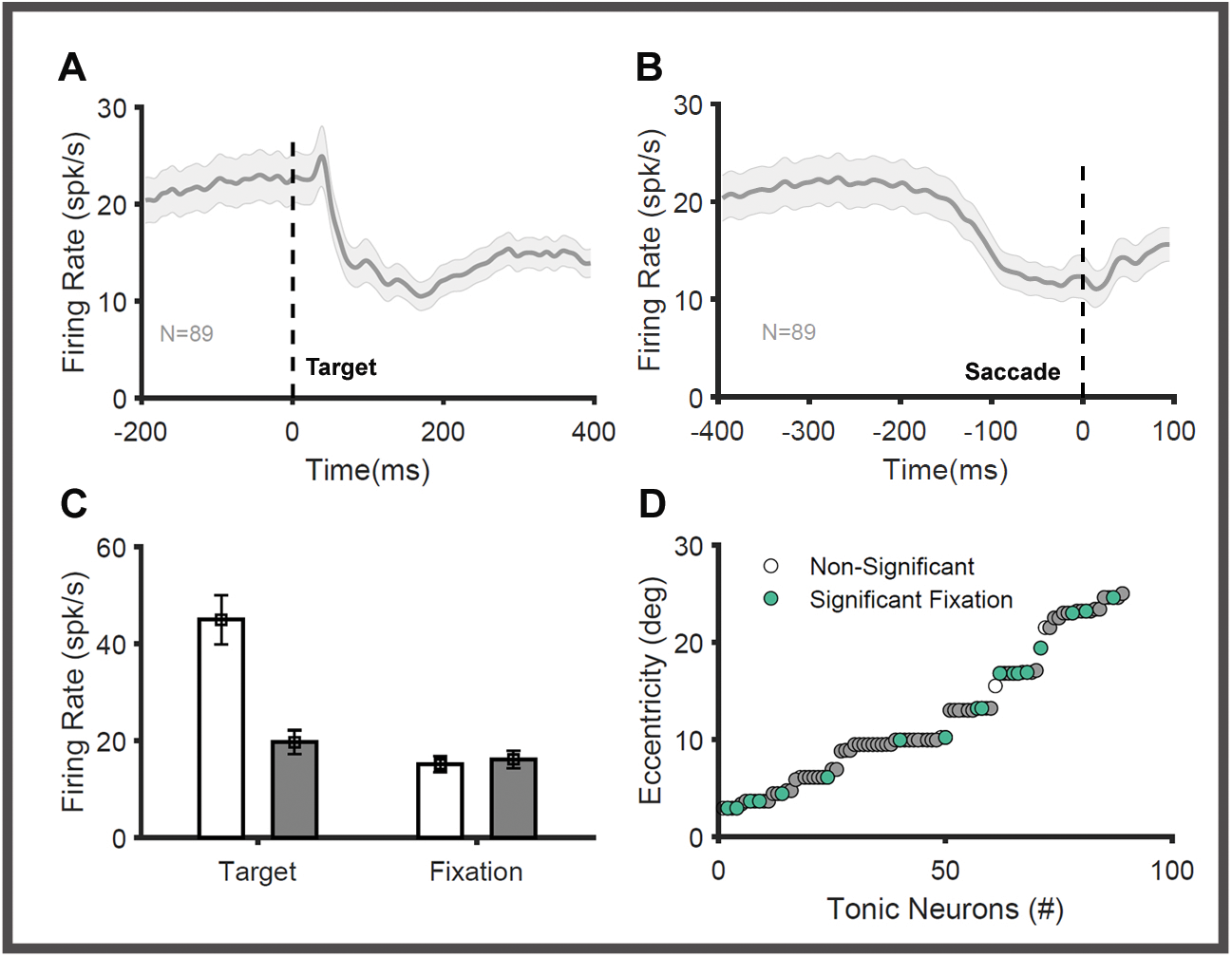
Properties of tonic neurons. The average response of SC tonic neurons when aligned on target onset **(A)** and saccade onset **(B)** is shown. **(C)** The mean firing rate from the population of tonic neurons in a 200 ms time interval before (white) and after (grey) target onset (left) and fixation onset (right) is shown as bar graphs. The error bars denote the standard error of means. **(D)** The eccentricity of the identified RFs of all the recorded tonic neurons (n=89) is shown. The green circles denote the tonic neurons, which had significant changes in activity in the 200 ms time interval before and after the onset of fixation.

Next, we also looked at the value responses of these tonic neurons. Unlike the other three groups of SC neurons (Fig-3A-C), there was no significant difference in their responses to good (12.4 spks/s) and bad objects (12.6 spks/s) when aligned on target onset (Fig-3D paired t-test p = 0.78). Similarly, no significant difference (paired t-test p = 0.69) was observed between good (11.56 spks/s) and bad (11.87spks/s) conditions during the pre-saccadic interval when the population activity of these neurons was aligned on saccade onset (Fig-3H paired t-test p = 0.70). Interestingly, this was not due to the absence of any significant modulation in these neurons. Instead, we found approximately equal proportions of positively (38%, n=34) and negatively modulated (43%, n=38) tonic neurons. The remaining 19% (n=17) of tonic neurons did not show any modulation in response to good and bad objects (Fig-3P).

### Identifying different epochs in SC neural responses

The three groups of SC neurons (visual, visuomotor, and motor neurons) play significant roles in transforming sensory visual stimuli into saccadic eye movements (Massot et al., 2019; Mays & Sparks, 1980; D. L. Sparks, 1986). The above data clearly shows that value-related modulation is observed in all three types of neurons, but it is unclear which epoch of the sensorimotor transformation is affected by this modulation. Does the value modulation in these neurons affect the visual processing or pre-saccadic motor stage? In this set of experiments, we used a visually guided saccade task, in which the visual and motor stages of sensorimotor transformations are seamlessly linked. Hence, to test this, we need to disentangle the visual and motor stages in the neural responses of these neurons, as done previously (Hanes & Schall, 1996; Thompson et al., 1996). If the neuronal activity in an epoch represents the visual processing, then it should exhibit no temporal variability when aligned on object onset. On the other hand, the motor processes will show temporal variability when aligned on object onset. If the neuronal activity in an epoch represents the pre-saccadic motor process, it should exhibit no temporal variability when aligned on the saccade onset; the visual process, on the other hand, will show temporal variability when aligned on the saccade onset. Using the above principle, we test if the value modulation seen in each neuron is temporally associated with the visual stage (onset of the object) or with the motor stages (i.e., start of the saccade motor plan) of the sensorimotor transformation.

To discriminate these stages of sensorimotor transformation in the neural activity of these SC neurons, we separated the trials into three groups based on saccade latency (e.g., short, medium, long). Next, we simulated neuronal population responses for each SC subtype using a bootstrapping method (SuppFig-2), which utilized the simultaneous recording of multiple neurons that we obtained using a linear array (see details in methods). It is well known that the activity of a single neuron does not directly control behavior. Instead, responses of multiple neurons are pooled together to direct a saccade in a trial. The bootstrapping method mimics this natural biological process unfolding in SC. In this analysis, we divided the neural responses recorded in different trials into three quantiles (short, medium, and long) based on saccade RT. We simulated the population response of a hypothetical trial belonging to a particular quantile by pooling single-trial responses of the same quantile from 25 random neurons. This process removed the variability in the single neuron response related to many non-specific factors such as object position, day-to-day variation in the animal’s motivation, etc. 15-35 trials were simulated for each quantile, and the mean firing rate and mean time of peak activity during different phases of the SC response were separately quantified. This bootstrap was repeated 10000 times to estimate the mean and the 95% confidence intervals.

The comparison of population activity simulated using the bootstrap procedure for the three RT quantiles identified the presence of different epochs with distinctive features. All three SC neuronal subtypes had an initial visual transient response that lasted for ∼80 ms. More importantly, the activity in this epoch peaked approximately (Fig:5B, F, J cyan line) at 42ms (range: 39-44ms) irrespective of the three RT groups for visual (Fig-5A), visuomotor (Fig-5E), and motor (Fig-5I) neurons. No temporal variability is associated with this peak when aligned on target onset, indicating that this epoch represents visual processing and is dubbed as early visual response (E_VIS_).

**Figure 5:**
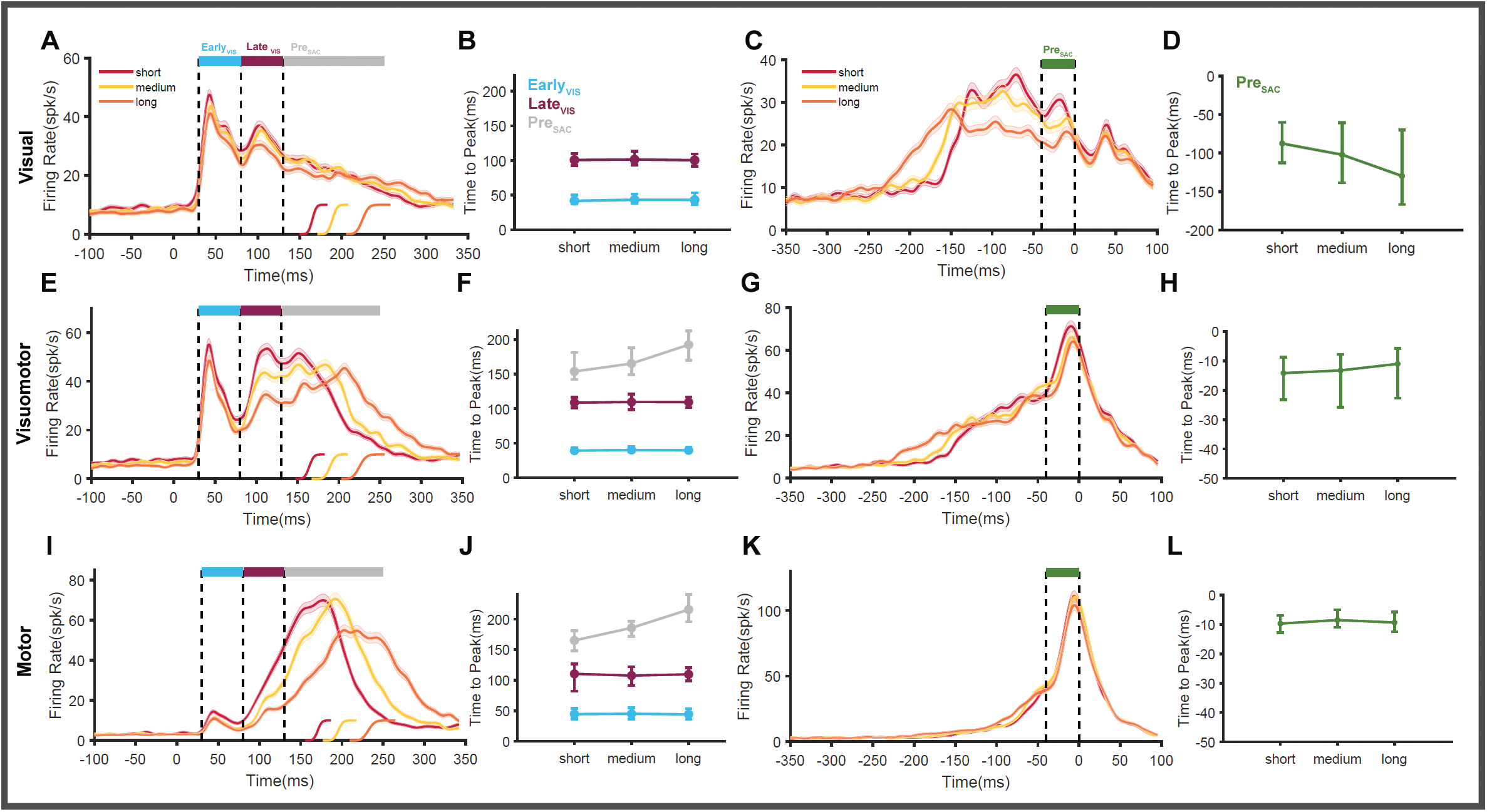
Identifying different phases of SC neuronal response. Simulated population activity of SC neurons generated by bootstrapping individual neuron responses separated by short (red), medium (yellow), and long (orange) RT trials are plotted. The activity is aligned on target onset and plotted separately for Visual **(A),** Visuomotor **(E),** and Motor **(I)** neurons. The vertical dashed lines denote the three phases of SC response: 1) early visual (E_VIS_) component (cyan) between 30-80ms, 2) late visual (L_VIS_) component (maroon) between 80-130ms, 3) tertiary pre-saccadic (PRE_SAC_) component (grey) beyond 130ms. The saccade RTs of trials belonging to the short (red), medium (yellow), and long (orange) groups are shown separately at the bottom of the graphs. The average time at which firing activity peaked in each RT quantile is shown separately for the E_VIS_ (cyan), L_VIS_ (maroon), and PRE_SAC_ components (grey) for Visual **(B),** Visuomotor **(F),** and Motor **(J)** neurons. Simulated population activity of SC neurons is aligned on saccade onset and plotted separately for visual **(C),** visuomotor **(G),** and motor **(K)** neurons. The vertical dashed lines denote the pre-saccadic phase of SC response:-40 to 0ms (green). The average time at which firing activity peaked in the pre-saccadic phase for each RT quantile is shown separately for visual **(D),** visuomotor **(H),** and motor **(L)** neurons. The error bars denote the 95% confidence interval in the peak time measured across the 10000 bootstrapped repeats.

Following this initial epoch, the activity in visual and visuomotor neurons increased again, resulting in a second peak within the interval of 80-130ms. Although the saccade latency (short, medium, long) strongly affected the level of activity in this epoch, the activity reached the peak approximately at 106 ms in visual (101 ms) and visuomotor neurons (109 ms) and motor neurons (108-110 ms) (maroon line in Fig-5B, F, J). Since there is no variability in the peak time when aligned on target onset, the activity in this second epoch is also related to visual processing and henceforth called the late visual response (L_VIS_).

After the L_VIS_ response (> 130ms), visual, visuomotor, and motor neurons continued to be active. Unlike early and late visual responses, the activity peaked in this epoch at different times, ordered by the difference in the saccade latencies of short, medium, and long RT groups. This distinction is clear for visuomotor (Fig-5E) and motor (Fig-5I) neurons (grey line in Fig-5F, 5J) but not so well for visual neurons (Fig-5A). This indicates that the activity after the L_VIS_ epoch may be temporally correlated with the saccade onset, representing the pre-saccadic motor process. Indeed, when the same activities are aligned on the saccade onset, their peaks are aligned on the saccade onset in the visuomotor (Fig-5G) and motor (Fig-5K) neurons. The timing of the peak activity in this third epoch was found to be ∼11ms before saccade onsets (Fig-5H, L) indicating that the activity represents the pre-saccadic (Pre_SAC_) motor response. In the case of visual neurons (Fig-5C), the peak activity was not aligned on saccade onset, but they varied systematically (Fig-5D) with respect to the RT groups, suggesting that these neurons encode only visual processes.

These data suggest that SC neurons (except tonic neurons) tend to have three distinct epochs/phases of activity that occur sequentially. Consistent with the previous studies, we were able to identify that the first two epochs - E_VIS_ and L_VIS_ - represent the visual processing stages of sensorimotor transformation.(McPeek & Keller, 2002; Schall & Hanes, 1993) The third epoch represented motor planning and was called the pre-saccadic activity (Pre_SAC_). Next, we compared the results obtained from the above bootstrapping procedure to the analysis shown in Fig. 3. This comparison suggests that reward modulation observed in the SC neurons affects the late visual and pre-saccadic phases of neuronal response.

To test this more systematically, we applied the bootstrapping method separately for good and bad objects, as shown in Fig 6. The good and bad object responses in the short (Fig-6A,G,O), medium (Fig-6C,I,O), and long (Fig-6E,K,Q) RT trials are compared. The mean firing rate and the 95% confidence intervals in each of the three previously identified epochs (E_VIS_, L_VIS_, Pre_SAC_) are compared. In visual neurons (Fig-6 B) and visuomotor neurons (Fig-6H), we found that E_VIS_ responses were not strongly affected by the object value (red vs. blue). In the motor neurons (Fig-6N), where the initial visual response is very weak, we found only negligible differences in the activity between good and bad conditions. In contrast to the E_VIS_ phase, the activity during the L_VIS_ phase was stronger for good objects than bad objects in all three subtypes of neurons: visual neurons (below the maroon line in Fig-6A,C,E and Fig-6D), visuomotor neurons (below the maroon line in Fig-6G,I,K, and Fig-6J) and motor neurons (below the maroon line in Fig-6M,0,Q and Fig-6P). In the Pre_SAC_ phase, where the activity represented the motor process in SC, we found that good objects elicited stronger responses than bad objects in the case of visuomotor (below the grey line in Fig-6G,I,K) and motor (below the grey line in Fig-6M,O,Q) neurons. The Pre_SAC_ activity was different based on object value (good object > bad object Fig-6L,R) in all saccade RT groups (short, medium, and long). Even though there is no clear pre-saccadic buildup of activity in the case of visual neurons (below the grey line in Fig-6A,C,E), there are clear differences between good and bad conditions in all three RT quantiles (Fig-6F). This analysis clearly indicates that reward modulation affects both the visual (L_VIS_) and the motor (Pre_SAC_) stages of SC neuronal response.

**Figure 6:**
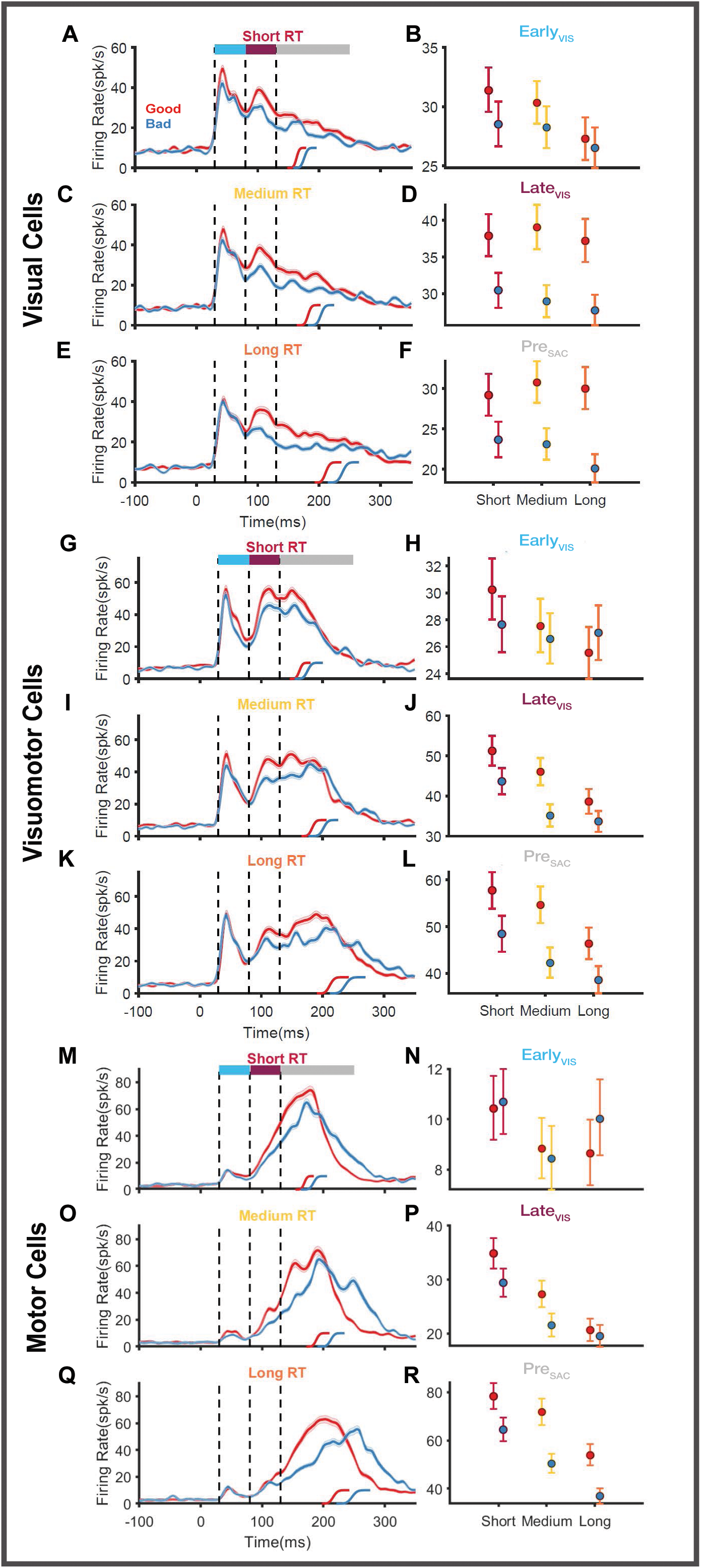
Effect of value on different phases of SC response. Simulated mean responses of visual neurons to good (red) and bad (blue) objects in the RF generated by bootstrapping individual neuron responses separately for short **(A)**, medium **(C)**, and long **(E)** RT trials are plotted. The shaded envelopes denote the standard deviation in the spike density functions measured across the 10000 repeats of the simulations. The saccade RTs of good and bad object trials are shown separately at the bottom of the graphs. The vertical dashed lines denote the different epochs identified in the responses of each SC subtype. The line graphs plot the mean firing rate and the 95% confidence intervals for the good (red) and bad (blue) object responses during the E_VIS_ **(B)**, L_VIS_ **(D)**, and PRE_SAC_ **(E)** phases of the response. The value effect in each RT quantile (short, medium, long) is separately shown in each graph. Simulated population responses of visuomotor neurons **(G-L)** and motor neurons **(M-R)** generated by bootstrapping individual neuron responses separated by short (top), medium (middle), and long (bottom) RT trials are plotted. The rest of the convention is the same as in panels A-G.

### Function of Tonic neurons

Previous analysis has shown that tonic neurons as a population are not clearly modulated by reward since they have equal proportions of positively and negatively modulated neurons. To identify the role of these neurons in sensorimotor transformation, we simulated the population activity of tonic neurons that gave rise to the short, medium, and long RT trials. The simulated population activity of tonic neurons (Fig-7) in each RT quantile is not significantly different from each other and peaked ∼ 49 ms [47 to 50ms] in the initial E_VIS_ epoch. In the following L_VIS_ epoch, the tonic neuron activity in all RT quantiles peaked ∼108ms. In the pre-saccadic phase, the activity of the tonic neurons reaches its minimum value (trough) prior to the onset of the saccades in an orderly fashion. The activity related to the short RT trials reaches the minimum first (157ms), followed by the medium RT trials (172ms), and then the longer RT trials (191ms, Fig-7B). This shows that the rate of inhibition in tonic neurons is correlated with the onset of saccadic eye movements. The temporal variability in reaching the lowest firing when aligned on target onset suggests that this activity is indeed related to motor processing (Fig-7A). When aligned on saccade onset, the variability in the motor processes is eliminated (Fig-7C, D) with the activity for all RT quantiles reaching the trough 33ms prior [-29 to-38ms] to the saccade onset. Given this, we suggest that the inhibitory activity in tonic neurons may directly contribute to the onset of saccadic eye movements.

**Figure 7:**
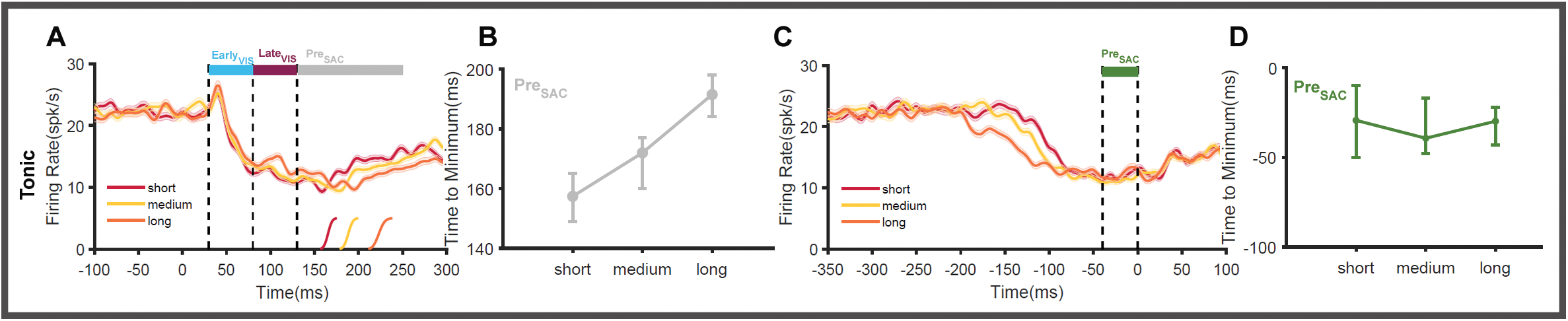
Role of Tonic neurons in saccade generation. Simulated population activity of tonic neurons generated by bootstrapping individual neuron responses separated by short (red), medium (yellow), and long (orange) RT trials are plotted aligned on target **(A)** and saccade **(C)** onset. The time at which the activity reached its minimum firing rate is calculated for each RT quantile and plotted when the activity is aligned on the target **(B)** and saccade onset **(D)**. The error bars denote the 5% confidence interval in the minimum time calculated across the 10000 repeats of the simulations.

### Value-modulated SC neurons control saccade behavior

According to the data so far (based on bootstrapping method), we found that the activity of SC neurons (visual, visuomotor, motor) may be divided into three phases: 1) Early visual response (E_VIS_), which is not modulated by object value 2) Late visual response (L_VIS_), which is strongly modulated by object value (good > bad) 3) Pre-saccadic activity (Pre_SAC_), which is slightly modulated by object value (good > bad). It is also known that SC population responses have a systematic relationship to the upcoming saccade in that higher firing rates accompany shorter RT and higher PV (Dorris et al., 1997; Waitzman et al., 1991). Taking these together, we hypothesize that SC neurons may act as an interface to functionally associate the object value with the behavioral enhancement of saccades to rewarding objects. This leads to the prediction that the degree of reward modulation found in each neuron will correlate with the strength of that same neuron’s influence on saccade behavior.

To test this prediction, we used three metrics that quantified the different influences on SC neural activity. Reward modulation in each neuron is parameterized by quantifying the difference between the average firing rate of good and bad objects during the time epoch 80-160ms (Value Index). Next, we quantified the impact of individual SC neurons on saccade behavior by calculating Pearson’s correlation between the firing rates and RT / PV separately. We defined the RT Index as the correlation between firing activity in the interval (80-160ms after target onset) and the corresponding RT in that trial for each SC neuronal subtype (SuppFig-3A-D). Similarly, we used the correlation between firing rate in the same epoch (80-160ms) and peak velocity (PV Index) to quantify individual neurons’ influence on the execution of the upcoming saccade separately for each SC neuronal subtype (SuppFig-3E-H). As expected, higher neural activities were associated with shorter RT (negative correlation SuppFig-3A-D) and higher PV (positive correlation SuppFig-3E-H). The correlation was calculated in three separate epochs:-100 to 0 ms, 0 to 80 ms, and 80-160 ms. Interestingly, the number of neurons with a significant correlation between firing rate and RT increased in epochs closer to saccade initiation, reflecting the evolution of the motor plan in SC (SuppFig-3I-L). As expected, a higher number of motor neurons (46%) have their firing activity significantly correlated with RT in the time interval 80-160ms compared to visual (22%), visuomotor (35%), or tonic (21%) neurons. The number of neurons with firing activity correlated with PV (8% averaged across 4 subtypes) is significantly lower than the number of neurons (20% averaged across 4 subtypes) modulated by RT.

We found significant correlations between value index and behavioral indices across neurons (Fig-8), confirming our prediction that the degree of reward modulation found in each neuron will correlate with the strength of that same neuron’s influence on saccade behavior. In visual neurons, the correlation was-0.312 (p<0.001 Fig-8A) between value and RT indices, and +0.216 (p<0.01, Fig-8B) between value and PV indices. The correlations between RT and value indices dramatically increased for visuomotor (-0.505, p<0.001 Fig-8C) and motor neurons (-0.579 p<0.001 Fig-8E). In contrast, the correlation between PV and value indices did not change in the case of motor neurons (+0.211, p<0.001 Fig-8D) and was not significant in the case of visuomotor neurons (+0.094, p=0.38 Fig-8F). In the tonic neurons, the correlation between value and RT indexes was-0.446 (p<0.001, Fig-8G), and +0.374 (p=0.003 Fig-8H) for correlation between value and PV indices. This analysis suggests that the heightened SC response to good objects can translate into shorter RT and higher PV. Thus, SC neurons act as a bridge between the value-processing circuits in the brain and the motor circuits that control saccades.

**Figure 8:**
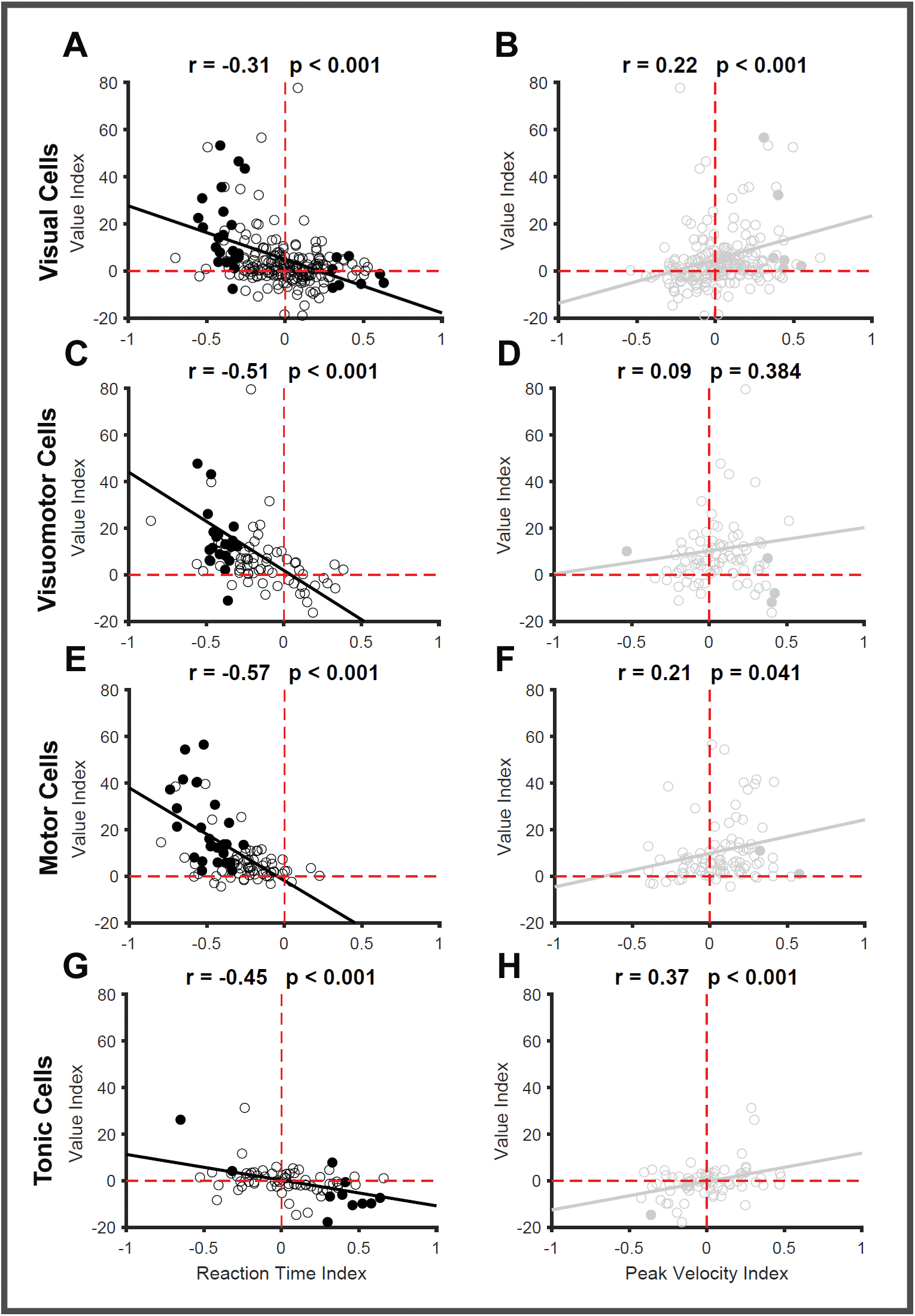
Relationship between reward and behavior modulation. Scatterplot showing the relation between reaction time index (correlation between firing rate and reaction time) and reward modulation (the difference in firing rate between good and bad objects in the time interval 80-160ms). Relations are assessed separately for visual **(A),** visuomotor **(C),** motor **(E),** and tonic **(G)** neurons. Each circle represents a single SC neuron. The best-fit regression line (black dashed line) is shown with a negative slope in all cases. Filled circles denote the neurons that showed significant reward and RT modulation indices. Scatterplots of peak velocity index (correlation between firing rate and peak velocity) vs. reward modulation (the difference in firing rate between good and bad objects in the time interval 80-160ms) are shown for visual **(B),** visuomotor **(D),** motor **(F),** and tonic **(H)** neurons. The best-fit regression lines have positive slopes in all cases. Filled circles denote the neurons that showed significant reward and PV modulation indices.

Next, we examined the distribution of the reward, RT, and PV modulated cells by plotting their cumulative probability against the indices used to quantify the extent of modulation (Fig-9A-C). The value modulation was quantified using AUROC, which spans from 0-1. We found that 70-80% of the recorded neurons in each subtype of the SC neuron were reward-modulated. More interestingly, the majority of visual (68%), visuomotor (84%), and motor (97%) neurons are positively modulated, while only 46% of the tonic neurons show the same positive modulation (Fig-9A). RT modulation was seen in 31% (averaged across the 4 subtypes) of the recorded neurons and was significantly lower than the proportion of reward-modulated cells (74%) seen in the recorded population of neurons. The majority of the RT-modulated cells in the visual (67%), visuomotor (100%), and motor (100%) neurons had higher firing rates when saccades were initiated with short latencies (negative correlation Fig-9B). On the other hand, only 31% of the RT-modulated tonic neurons have their firing rates negatively correlated to reaction time, while the remaining 69% of cells are positively correlated.

**Figure 9:**
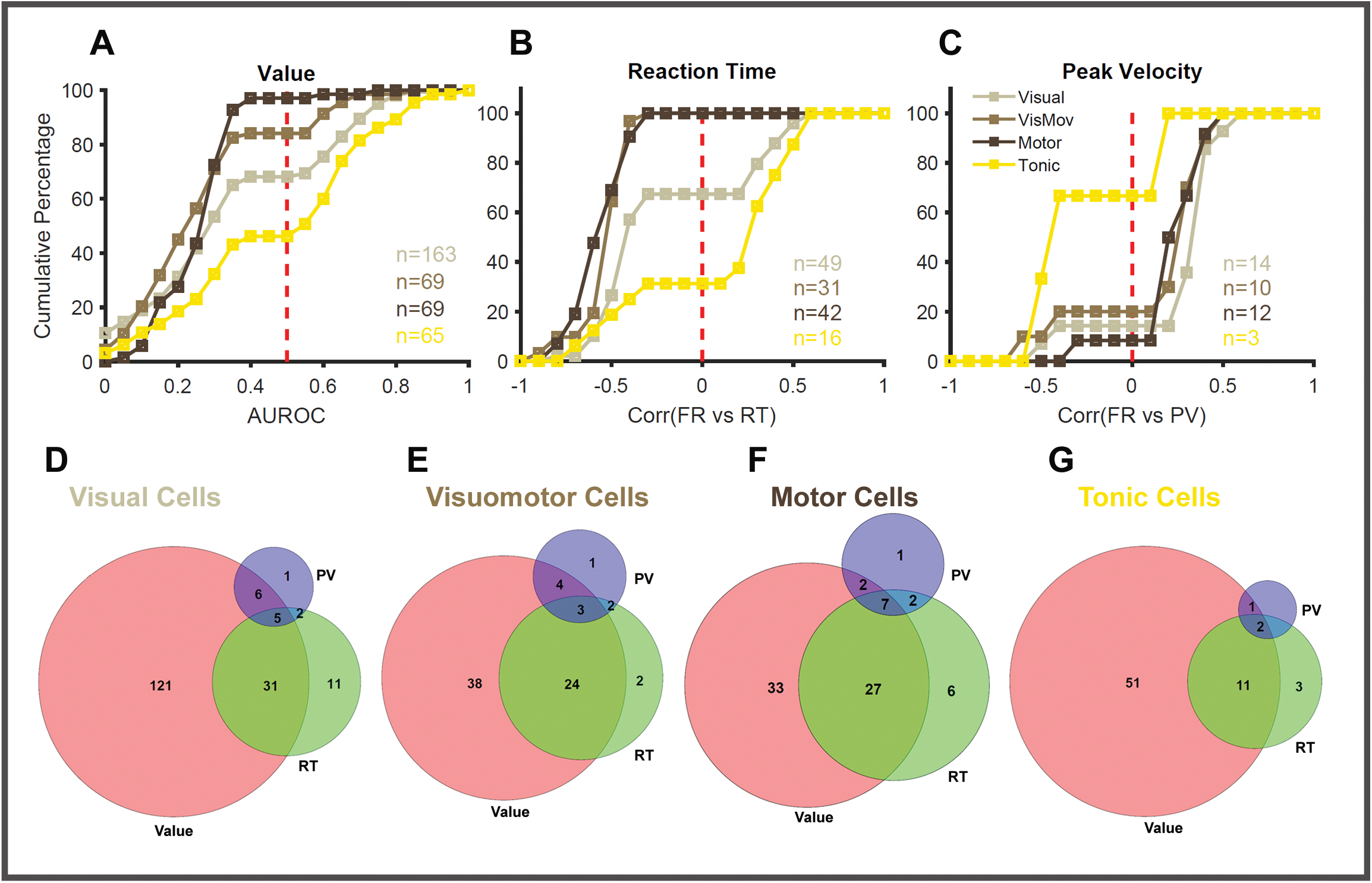
Distribution of modulated SC neurons **(A)** Cumulative plot showing the number of neurons that are modulated by reward from visual (grey), visuomotor (brown), motor (black), and tonic (yellow) neurons in SC. The reward modulation is assessed using AUROC. The neurons with AUROC values from 0 to 0.5 denote positively modulated cells, and those from 0.5 to 1 denote negatively modulated neurons. The total number of neurons from each SC subtype modulated by reward is also shown in their corresponding colors. **(B)** Cumulative plot showing the number of neurons that are modulated by RT from visual (grey), visuomotor (brown), motor (black), and tonic (yellow) neurons in SC. The RT modulation is quantified as the Pearson’s correlation between firing rates in the time interval 80-160ms and the corresponding trial RT. The neurons with RT indices ranging from-1 to 0 denote negatively modulated cells, and those from 0 to 1 denote positively modulated neurons. The total number of neurons from each SC subtype that are RT modulated is also shown in their corresponding colors. **(C)** Cumulative plot showing the number of neurons that are modulated by PV from visual (grey), visuomotor (brown), motor (black), and tonic (yellow) neurons in SC. The PV modulation is quantified as Pearson’s correlation between firing rates in the time interval 80-160ms and the PV of the corresponding trial. The neurons with PV indices ranging from-1 to 0 denote negatively modulated cells, and those from 0 to 1 denote positively modulated neurons. The total number of neurons from each SC subtype that are modulated by PV is also shown in their corresponding colors. Venn diagram showing the overlap between reward (pink), RT (green), and PV (blue) modulated neurons calculated separately for visual **(D)**, visuomotor **(E)**, motor **(F)**, and tonic **(G)** neurons. The intersection area where all three circles meet denotes the neurons that are modulated by all three factors. The intersection area where two circles meet denote neurons that are modulated by reward/RT, reward/PV, and RT/PV. The area of the circle that does not intersect with the others denotes neurons modulated by only one factor. The number of neurons belonging to each of the above groups is also given.

In contrast, PV modulation is seen only in a minority of SC cells (9% averaged across 4 subtypes) compared to the value (74%) and RT (31%) modulated cells. The majority of these PV-modulated visual (86%), visuomotor (80%), and motor (92%) neurons have their firing rates positively correlated with the PV of the upcoming saccade (Fig-9C). As before, only 33% of the tonic neurons had firing rates that were positively correlated to PV. This analysis shows that there are many more reward-modulated neurons in SC, out of which a smaller proportion of cells play a significant role in converting the reward signal into behvaior - saccadic vigor.

We also looked at the overlap between the reward, RT, and PV-modulated cells separately in each functional subtype of SC neurons (Fig-9D-G). Similar patterns of overlap were seen across all four subtypes of SC neurons. Interestingly, only a very small proportion of visual (2%), visuomotor (3%), motor (7%), or tonic (2%) neurons were modulated by all three factors simultaneously. Among the visual and tonic neurons, 14% of the cells were modulated by both reward and RT. The proportion of such dual (reward/RT)-modulated neurons increased in the case of visuomotor (27%) and motor (30%) neurons. The number of dual (reward/PV)-modulated neurons was similar across the 4 subtypes of SC neurons and was much smaller in proportion (3%) compared to the other type of dual (reward/RT) modulated neurons. This analysis suggests that PV and RT modulated cells are largely non-overlapping subgroups within the group of value-modulated SC neurons.

### Value signals interface with the local circuits in the superior colliculus

The analysis so far showed clearly that the different subtypes of SC neurons, despite their different functional roles, respond similarly to the reward associated with objects. In the visual, visuomotor, and motor neurons, the activity differences appeared only during the L_VIS_ and the Pre_SAC_ phases of the response, while the activity during the E_VIS_ phase did not distinguish between good and bad objects. The delayed onsets of reward modulations in these three types of neurons suggest that the neural activity in SC evolves over time: E_VIS_ in response to the appearance of visual objects in the RF and later L_VIS_ in response to the value of the object. To test this, we quantified the time when the activity profiles of good and bad objects diverged significantly from each other, i.e., the onset of reward modulation (See methods). To visualize the reward modulation (Supplementary Fig-4) more clearly, the difference between the activity of good and bad objects (SDF_good_ – SDF_bad_) was calculated and plotted. On average, the visual (SFig-4A), visuomotor (SFig-4B), and motor neuron (SFig-4C) show a positive value modulation (higher activation for good objects). Since tonic neurons do not show any clear value responses as a population, they were not included in this analysis. The time of reward modulation onset was calculated from the averaged activity difference between good and bad conditions. We observed that the value information appeared in the visual (86 ms) and visuomotor neurons (87 ms) approximately at the same time, while it appeared in the motor neurons (100 ms) about 10 ms later (SFig-4A-C). Next, we repeated the same procedure in individual neurons and got a distribution of reward modulation onsets separately for each functional subtype of SC neurons. We did not observe any significant difference between the distributions of onset values in the three classes of SC neurons (p=0.167, Kruskal-Wallis Test). On average, the effect of reward modulation became significant at 98 ms in visual neurons (SFig-4D), 95 ms in visuomotor neurons (SFig-4E), and 113 ms in the motor neurons (SFig-4F). This shows that reward-related afferent inputs from other cortical or subcortical sources interface with the three functional subtypes of SC neurons, thereby modulating the vigor of the upcoming saccade to high-valued objects.

## DISCUSSION

We investigated how the value-sensitive inputs from other brain regions interact with different subtypes of neurons in the SC that form the visuomotor circuit and modulate saccadic behavior. Using linear arrays, we assessed reward modulation in four different subtypes of SC neurons while the animals were engaged in a visually guided saccade task, which is attuned to the natural behavior of the animal. SC is an important hub of sensorimotor transformation; hence, the neurons are involved in both the visual processing of the target and the motor preparation of saccades. A bootstrapping method was used to simulate the population activity of different neuronal subtypes and identify the various phases of sensorimotor transformation in the response profiles of SC neurons. We identified three distinct phases in SC responses: an early visual phase agnostic to value, a late visual phase, and a tertiary presaccadic motor phase that are modulated by the value of the target present in the RF of the neurons. Three out of four SC neuronal subtypes respond with higher firing rates to high-valued good objects than to low-valued bad objects during the late visual and the pre-saccadic epochs of activity, which resulted in an enhanced saccadic response toward good objects. Additionally, we identified a linear relationship between the extent of an individual neuron’s value modulation and the influence of the same neuron on saccade RT and PV. The greater the reward modulation, the greater the cell’s impact on saccade behavior (Fig-8). In conclusion, this study comprehensively assessed the effect of reward on different functional subtypes of SC neurons, suggesting that they act as a bridge between the circuits that process value information and that control motor actions.

### Value response in SC neurons

In this study, we observed that saccades directed to high-valued objects are initiated and executed faster than those directed to low-valued objects. This observation is consistent with other studies in humans (Milstein & Dorris, 2007; Reppert et al., 2015; Sedaghat-Nejad et al., 2019) as well as NHPs (L. L. Chen et al., 2013; Takikawa et al., 2002) that reported similar value-driven saccadic invigoration.

We observed higher firing rates in three functional subtypes of SC neurons when they directed their saccades to high-valued objects in the RF of these neurons in the time interval 80-160ms after target onset. Interestingly, previous studies that attempted to address reward modulation in SC neurons reported different responses (Ikeda & Hikosaka, 2003, 2007). In those studies, visual neurons showed two types of significant modulations in response to reward. 1) bias modulation, in that the activity of neurons before the target onset (baseline) showed a ramping of activity well correlated with the value of the upcoming saccades; 2) gain modulation, in that the initial transient response (E_VIS_ here) of SC neurons showed differential activation based on the value of upcoming saccade targets (Ikeda & Hikosaka, 2003). In addition, the value modulation observed in the motor neurons have two distnict temporal profiles: 1) positive reward coding neurons exhibited higher activation throughout the trial, starting prior to the presentation of the cue till the execution of the saccade 2) negative reward coding neurons exhibited higher activation to low value largely restricted to the pre-saccadic buildup phase (Ikeda & Hikosaka, 2007). In contrast, this study found that the value responses observed in three subtypes of visual, visuomotor, and motor SC neurons shared some common features. The baseline activity and the E_VIS_ phases of the response did not differentiate between objects’ values, while the L_VIS_ and Pre_SAC_ phases are significantly modulated in all three neuronal subtypes (Fig-3,6). In addition, both the positively and negatively modulated neurons reported here showed opposite firing patterns in response to value, but they had similar temporal profiles of activation.

We hypothesize that this variation seen in the responses of SC neurons may arise from the differences in tasks used in these studies. Here, the value was associated with the identity of an object through repeated and consistent exposure for more than ten days, giving rise to long-term object value. In the earlier two studies, an asymmetrically rewarded (1-DR) task was used in which one target position on the screen was associated with a higher reward while the rest of the positions gave low rewards. The reward assigned to the location flipped after the end of a block (∼30 trials), giving rise to the short-term position-value association. Given this difference in task design, there are two plausible explanations.

(1) It is likely that SC neurons encode the positional and object values differently. The positional value response appears in SC visual neurons even before target onset in“bias-type” neurons, whereas in “gain-type” neurons, value modulation appears concomitantly with the early visual transient (Ikeda & Hikosaka, 2003). In contrast, the modulation due to object value appeared in SC neurons in the interval 80-160ms after target onset. We intend to test this in a follow-up study by recording the same pool of neurons while the animal performs the two different tasks.
(2) Previous studies have shown two parallel circuits in basal ganglia (Hikosaka et al., 2019; H. F. Kim & Hikosaka, 2015) that separately mediate memory for long-term stable values (such as the stable object task used in the current study) and memory for short-term flexible values (such as the 1-DR task). The flexible circuit originates from the head of the caudate nucleus and projects to the rostral ventral medial (rvm) part of SNr. On the other hand, the stable circuit originates from the tail of the caudate nucleus and projects to the caudal dorsal lateral (cdl) part of the SNr (H. F. Kim & Hikosaka, 2013; Yamamoto et al., 2013; Yasuda & Hikosaka, 2015). Both the circuits from rvmSNr and cdlSNr converge onto SC and modulate behavior. The differences in the value response of SC neurons between the two studies may arise from the differential activation of these parallel circuits in BG triggered by the long-term stable vs. short-term flexible values involved in the tasks used.

The higher activation seen in these neurons might be related to the value of the expected reward or to the degree of motivation that the animal has to acquire the reward. Prior experiments (Roesch & Olson, 2003, 2004) were conducted in which the monkeys were trained to saccade towards objects that provided different volumes of juice reward (positive outcome) and objects that necessitated a long wait time before the next trial (negative outcome). In this task, the degree of motivation was controlled independently by the magnitude of the reward promised in the event of success and the magnitude of the penalty threatened in the event of failure. The large reward and long wait time cues have high behavioral relevance, and the neurons sensitive to motivation will have similar activation profiles. In contrast, the neurons sensitive to the expected value will have distinct activation profiles since the animals clearly prefer the positive outcome. Unfortunately, the task design used here does not allow us to distinguish between these alternatives. These studies have further shown that neuronal representations of expected value, independent of the actions, are seen in prefrontal regions, while the more posterior regions in the saccade network are strongly influenced by the motivation to acquire the reward and the ensuing motor preparation. (Roesch & Olson, 2007) The neural activity in SC is also known to be related to motor output; hence, the reward modulation we observed in SC neurons is likely related to the motivation the animal had to acquire the high-valued object. The strong relationship observed here between the reward modulation and the saccade parameters strongly favors this possibility.

Anticipation of a high-valued reward leads to stronger motivation, as well as greater arousal and attention. Interestingly, previous studies have shown that the activity of SC neurons is strongly influenced by such factors (Fecteau et al., 2004; Herman & Krauzlis, 2017; Ignashchenkova et al., 2004). For example, neurons have higher firing rates when an object appears in a location the subjects had covertly attended. The activity is lower when the object appears in the same location while the subject’s attention is directed elsewhere. High-valued objects, due to their higher behavioral relevance, tend to draw more attention compared to low-valued objects (Ghazizadeh et al., 2016b). Hence, it may have contributed to the activity modulation observed here. In other words, the reward modulation we observed here may also subserve the attentional mechanisms previously reported in SC or vice versa; however, given the task we employed here, it is not possible to distinguish the specific contribution of attention from reward.

Similarly, previous studies have also shown that SC neurons discriminate targets of the upcoming saccade from a distractor in its RF with higher firing rates during an oddball visual search task (B. Kim & Basso, 2008; R. Krauzlis & Dill, 2002). Visuomotor neurons showed a biphasic activation profile where the initial visual response did not discriminate the target from the distractor, while the second visual phase was strongly modulated to signal the targets (McPeek & Keller, 2002). Here, we find a similar biphasic response not only in the visuomotor but also in the visual and motor neurons. The same underlying mechanism of target discrimination may be utilized in the current task to discriminate between high and low-valued targets and manifest as the reward modulation during the late visual phase. Regardless of the underlying mechanism, the observed modulation in SC neurons with extrafoveal RFs would help the monkeys identify and saccade toward the high-valued object among a multitude of distracting objects presented in their peripheral vision (Ghazizadeh et al., 2016a).

In addition, the SC neurons in the superficial layers, which receive inputs from the early visual processing areas, are known to generate a saliency map of the external world (White, Berg, et al., 2017; White, Kan, et al., 2017). Hence, the objects that have higher luminance, contrast, or other such physical attributes will affect the firing rates of visual neurons. In this study, we used multiple sets of 8 fractal objects as targets, which were generated randomly with varying colors, contrast, shape, etc, and half of them were randomly assigned as high-valued objects. The animals were repeatedly exposed to these objects and their associated reward for more than 10 days, allowing them to learn the ecological salience of these fractal objects, which is verified using their gaze position under free viewing (Ghazizadeh et al., 2016b; H. F. Kim & Hikosaka, 2015). This procedure will nullify any systematic effects that low-level visual features of the objects have on the firing rate of SC neurons. Hence, the contribution of physical saliency to the observed reward modulation of visual neurons might be minimal.

### Source of value signals

The SC receives inputs from multiple regions of the brain, which are organized into different layers. The superficial input layers receive major inputs from the early visual areas in the cortex (Cerkevich et al., 2014; Jiang et al., 2023; May, 2006) and the retina (Cowey & Perry, 1980; Grünert et al., 2021; Rodieck & Watanabe, 1993), while the intermediate layers receive information from a variety of sources like the basal ganglia, frontal eye fields, supplementary eye fields, and lateral inter-parietal area (Inoue et al., 2015; Sommer & Wurtz, 2000; Wurtz et al., 2001). An early low-latency visual epoch is seen in all four subtypes of SC neurons, which is agnostic to the value of objects in the RF and is likely driven by the sensory inputs from multiple early visual regions. Conversely, the activity during the later visual epoch, i.e., 80-160ms after target onset, may be driven by inputs from other brain regions like the cdlSNr of the basal ganglia, FEF, LIP, etc. We have seen that activity in this later visual epoch, as well as the presaccadic motor epoch of SC response, is modulated by reward and may originate from these sources. More importantly, this shows that reward value does not indiscriminately alter sensory representations, but selectively modulates the later integrative stages of sensorimotor transformation.

One of the most prominent inputs to SC that actively modulates saccadic response is the output nucleus of BG, SNr, which tonically fires to inhibit SC neurons so that the animal can keep its eye fixated. Prior to a saccade, SNr neurons reduce their firing rates, disinhibit the SC, and facilitate movement initiation. (Hikosaka et al., 2006; Shires et al., 2010; Watanabe & Munoz, 2011). A previous study has shown that antidromic stimulation of visual, visuomotor, and saccadic SC neurons localized in different layers of SC can activate SNr neurons (Hikosaka & Wurtz, 1983). In addition, orthodromic stimulation of SNr neurons influenced the motor neurons in the SC (Liu & Basso, 2008). More importantly, the previous work from the lab showed that SNr neurons are phasically inhibited by a high-valued object, facilitating a saccade towards it. In contrast, the same neurons are phasically excited by a low value, which delays the saccade toward the bad object (Sato & Hikosaka, 2002; Yasuda et al., 2012). This long-term object-based value modulation of SNr activity is seen around 100 ms and is thought to be driven by visual input from the tail of the caudate nucleus (Yasuda & Hikosaka, 2015). Here, using a similar long-term stable value saccade task, we showed that the value modulation appeared in SC between 95-115ms after target onset in three subtypes of neurons (SupFig-4). This timing suggests that cdlSNr neurons may interface with visual, visuomotor, and motor neurons in SC, thereby inducing value responses quickly. This is supported by a recent study in which optogenetic activation of the CDt-cdlSNr pathway by laser applied in the SNr modulated the activity of SC visual neurons (Amita et al., 2020). Given this evidence, we speculate that value-sensitive output from the cdlSNr interfaces with all three subtypes of SC neurons localized in the different layers of SC and facilitates the observed behavioral modulation of saccades.

In addition to SNr, neurons in the frontal eye field also send saccade-related information to the SC (Matsumoto et al., 2018; Sommer & Wurtz, 2000) and are known to have neurons modulated by reward expectation (Glaser et al., 2016; Roesch & Olson, 2003). In a gambling task, they also showed persistently higher firing for high-valued objects even after the saccade, which was considered to be helpful in learning the reward association (X. Chen et al., 2020). The FEF neurons are known to have persistently higher firing for high-valued objects after the cue onset, and sometimes during their baseline periods (Ding & Hikosaka, 2006). Similar to the results shown here, all the 3 functional subtypes of neurons seen in the FEF were also affected by the value task, with visuomotor neurons having the highest number of modulated cells. In contrast, we have shown that the firing activity of ∼80% of visuomotor and motor neurons in SC is affected by the value of the objects. Similar neuronal responses to value have also been seen in other cortical areas like supplementary eye fields and lateral intraparietal area (Amador et al., 2000; Bendiksby & Platt, 2006; Coe et al., 2002; So & Stuphorn, 2010), which play a major role in generating saccades and send direct projections to the colliculus (Huerta & Kaas, 1990; Paré & Wurtz, 2001; Wurtz et al., 2001). Given the similarity of value responses in these different regions, it is challenging to speculate on the source of value inputs to the SC neurons. In addition, the tasks used in these studies were not comparable to the fractal-based visually guided saccade task used here. Moreover, the time of value onsets was also not quantified in these studies, making the comparison difficult. We intend to pursue this using more rigorous methodologies in future studies.

### Local circuits in SC for saccade invigoration

The superior colliculus has a well-defined laminar structure, with the different functional subtypes of neurons organized into specific layers. Previous research with SC slice recordings has shown local circuitry comprising connections between the different layers that facilitate the flow of activity in a structured manner from visual neurons in the superficial layers to the visuomotor neurons and motor neurons in the intermediate and deep layers (Isa, 2002; Isa et al., 1998; Isa & Saito, 2001). There are also feedback connections, both excitatory and inhibitory, from the motor neurons in the deep layers back to the superficial neurons (Ghitani et al., 2014; Phongphanphanee et al., 2011). This interacting local circuit in SC can have profound impacts on the incoming inputs. For example, a brief optogenetic stimulation of the cdlSNr-SC pathway (<20ms) had a prolonged effect (∼200ms) on SC responses (Amita et al., 2020). The brief disinhibition from cdlSNr may trigger the local excitatory interactions in the intermediate and deep layers of SC (P. Lee & Hall, 2006; Pettit et al., 1999), thereby enhancing the burst activity of motor neurons until a saccade is generated. Value inputs from other cortical and subcortical regions impinge on the intermediate layers of SC and may propagate through the local circuit to the other classes of neurons. Consistent with this idea, albeit statistically not significant, we found a trend that the value modulation first appeared in the visuomotor neurons at a latency of 95ms and then propagated to the visual (98ms) and motor neurons (113ms). Laminar probes allow us to track these local interactions between various cell types in the different layers of SC, and we intend to pursue this in the future.

We have shown that the L_VIS_ phase of SC neurons significantly differentiates between good and bad objects. The same activity also has a systematic relationship with the onset of saccades as well as the peak saccade velocities. Consistent with this result, previous studies have shown that motor neuron activity directly influences behavior by modulating saccade RT and peak velocity (Dorris et al., 1997; R. J. Krauzlis, 2003; Smalianchuk et al., 2018; Waitzman et al., 1991). In another report, the activity of visual neurons was modulated when stimuli of higher/lower intensities appeared in their RFs, resulting in the modulation of saccadic RT (Bell et al., 2006; Marino et al., 2012). In this study, we observed that all four subtypes of SC neurons have a systematic relationship to the initiation and execution of the saccade. The extent of an individual neuron’s modulation with respect to high and low reward is linearly related to the same neuron’s influence on saccade RT and PV. The greater the reward modulation, the greater the cell’s impact on saccade behavior. As expected, visuomotor and motor neurons exert relatively stronger influences on behavior than visual and tonic neurons. Hence, we were able to assess the relative contributions of different neuronal subtypes in controlling the value-based enhancement of saccadic behavior.

We also found that the neuronal activity of the majority of SC neurons (>70%) in each subtype is modulated by the reward that the animals received after the saccade. However, we found that only a small subset of reward-modulated neurons showed significant correlations between their firing rates and the RT(∼30%) and PV(∼8%) of the upcoming saccade (Fig-9). Interestingly, similar patterns were seen in other cortical regions, such as FEF, which is modulated by reward. It is possible that the non-saccade reward-modulated SC neurons send their outputs to modulate other effectors such as the neck and hand (Corneil et al., 2002; Stuphorn et al., 2000; Walton et al., 2007) and may facilitate the invigoration of the coordinated movements to high-valued objects, which is critical for animals foraging in natural environments. SC is an important hub that controls the escape behvaior – those innate, fast, reflexive responses seen in animals in aversive situations. The reward-modulated non-saccade neurons in SC may also project to the downstream regions like peri aqueductal grey or amygdala via pulvinar and control this escape behavior (Basso et al., 2021; Evans et al., 2018; Koller et al., 2019). In addition to output structures, these reward-modulated non-saccadic neurons may send ascending projections to other brain regions. There is a well-known direct projection from SC to substantia nigra compacta that is thought to generate a phasic response in dopamine neurons (Comoli et al., 2003; Redgrave et al., 2010). There is also evidence for the existence of disynaptic connections to the caudate nucleus through the centromedian and para fascicular nucleus of the thalamus from SC (Harting et al., 1980; Ichinohe & Shoumura, 1998; Sadikot et al., 1992). Through these pathways, the higher activity in SC neurons in response to rewarding stimuli may play a critical role in learning novel value associations.

### Tonic Neurons

In this study, we observed a class of SC cells that were named tonic neurons and had not been extensively studied previously. As the name suggests, these neurons are tonically active and are inhibited when a target appears in the cell’s RF in the contralateral visual field and remain inhibited till after the end of the saccade. These tonic neurons have activity profiles distinct from those of the foveal neurons in rostral SC, which briefly pause their tonic activity during the saccade execution (Hafed & Krauzlis, 2012; R. J. Krauzlis et al., 2000; Munoz & Wurtz, 1993). Unlike these foveal neurons, tonic neurons have RFs ranging from 4-25 degrees and are found mingled with other subtypes along the dorsoventral axis of SC. These neurons are also distinct from the visual tonic neurons found in SC, which show a brief pause in the delay period activity before the saccade onset, driven by the appearance of the central cue that indicates the nature of the upcoming movement (Li et al., 2006). Interestingly, we found that activity in the majority of the tonic neurons is unaffected by the onset of the central fixation cue. So, tonic neurons might be a novel class of neurons that has not been reported earlier.

We speculate that the tonic neurons may be part of the local inhibitory network, presumably, the GABAergic neurons reported previously in SC slice recordings, which help coordinate the flow of information between the different layers of SC. Fast-spiking GABAergic interneurons with different types of axonal morphologies are reported in the intermediate SC to provide intralaminar and interlaminar inhibition. These neurons play an important role in shaping the activations in the motor maps of SC and facilitate accurate saccades (P. Lee & Hall, 2006; Sooksawate et al., 2011). It has been shown that reducing the level of GABA-mediated inhibition in the local circuit can induce bursts of spikes in premotor intermediate-layer neurons with minimal excitation in the superficial visual layers (Kaneda et al., 2008). Hence, these local inhibitory neurons may modulate the firing of premotor neurons in the intermediate layers to facilitate faster saccades or further inhibit the non-target motor neurons, thereby sharpening the activity in the SC motor maps to facilitate accurate saccades. Consistent with this idea, we found that when the activity of tonic neurons reached their lowest activity state earlier, it resulted in faster saccades RTs, suggesting a potential disinhibitory influence. We suggest that when there are no objects in the RF, higher baseline firing in the tonic neurons may inhibit the interaction between the other classes of SC neurons. Hence, they may play a significant role in enabling stable fixation, which is very important for perception, attention, and social interaction. When an object (good or bad) appears, tonic neurons lower their firing activity (Fig-4A,B), which would disinhibit visual, visuomotor, and motor neurons, thereby facilitating saccades. We speculate that this mechanism of disinhibition by tonic neurons is particularly relevant in the case of bad objects, where a saccade has to be generated towards the object despite lower activation in the visual, visuomotor, and motor neurons. We have also seen equal proportions of positive and negatively modulated tonic neurons, suggesting that they may have a complex interaction with other subtypes of SC neurons to modulate saccade behvaior.

Taken together, this study provides a comprehensive analysis of the various functional cell types of SC neurons and their role in invigorating saccadic responses to high-valued visual objects in the environment. These results illustrate a circuit-level mechanism by which a subcortical sensorimotor hub integrates motivational information into the visuomotor planning to facilitate value-guided behvaior.

## Acknowledgments

We thank Hikosaka lab members for their critical, in-depth discussion of these results. We also thank A.M. Nichols, D. Yochelson, G. Tansey, D. Parker, I. Bruna, A. Lopez, and H. Warnock for their technical assistance. We acknowledge Dr. Leor Katz’s help in setting up the spike-sorting pipeline used in this study. We also thank Dr. Richard Krauzlis for the support provided while performing experiments and drafting this manuscript. This research was supported by the Intramural Research Program of the NIH, National Eye Institute.

## Author contribution

AG and OH designed these experiments. AG performed animal training, surgeries, and neural recordings. AG also performed data analysis and prepared the figures. AG, and OH drafted the manuscript. AG edited and revised the manuscript.

## Data and Code Availability

The data and the matlab scripts that support the findings of this study are available from the corresponding author upon reasonable request.

## MATERIALS AND METHODS

### Animals and Surgical Procedures

Two non-human primate (macaca mulatta) males with weights of 9kgs (DWN) and 13kgs (BLY-were used in this experiment. All animal care and experimental procedures were approved by the National Eye Institute Animal Care and Use Committee and followed the Public Health Service Policy on the Humane Care and Use of Laboratory Animals. Survival surgeries were performed under sterile conditions and isoflurane gas anesthesia. A head holder was implanted at the stereotaxic zero to restrict the head movements of the animal. A rectangular recording chamber was implanted posterior to the head post at an angle of 40 degrees off vertical (Fig-1C). This orientation of the recording chamber ensured that the receptive fields (RF) of the different neurons recorded simultaneously from the different layers of the SC mostly overlapped.

### Recording Setup and Procedure

The monkeys performed a visually guided saccade task (Fig-1A) that was controlled by a custom real-time software, BLIP (www.simonhong.com). The visual stimuli (Fig-1B) used in the task were fractal objects (Yamamoto et al., 2013) (average size ∼7° X 7°, range 5-10°) with several randomly determined features (amplitude, colors, and shapes) created uniquely for these monkeys. We used four distinct fractal sets, each consisting of 8 objects, for each monkey. The objects were randomly assigned to high and low-valued groups. This was done to avoid differences in the physical properties of the stimuli systematically affecting our results. Eye position was tracked at 1 kHz using an infrared eye-tracking system (EyeLink1000, SR Research). We used a 24-channel, linear electrode array (V probe – Plexon Inc), with inter-electrode distances (50 microns/100 microns), to record the neural activity from different layers of the SC simultaneously. The signals from the 24 channels of the V-probe were filtered, amplified, and digitized using a 64-channel OMNIPLEX-D data acquisition system (Plexon Inc) at a sampling frequency of 40 kHz.

In a typical experimental session, we chose a grid location that targets the SC based on MRI images (Fig-1C), and a 24-channel v-probe was then lowered into the brain using a manual hydraulic manipulator (MO-97A Narishige Int’l. USA). We encountered SC neurons, typically at a depth of 40mm from the bottom of the grid, which are characterized by visual neurons with low latency phasic response when a target is presented in a specific location of the visual field. After identifying the dorsal-most layer of SC, the probe was lowered further until all 24 channels on the probe were in contact with different layers of the SC. A v-probe with a 100-micron inter-electrode distance spans 2400 microns of SC and can typically record multiple functional subtypes of neurons across the different layers simultaneously (Fig-1C).

We used a passive viewing task to map the neurons’ receptive fields. The monkeys were trained to fixate at the center of the screen while objects were flashed at different visual field locations. The stimuli were presented along eight radial angles, starting at zero (horizontal) at 45-degree increments and with amplitudes ranging from 5 to 25 degrees at 5-degree increments. The screen location that elicited high firing in the most number of channels along the dorsoventral axis of SC was identified as the RF and used in the subsequent behavioral tasks. This approach ensured that most of the neurons we recorded had some task-specific responses, even if the targets were not placed in their ideal fields due to the inability to finely tune and identify RFs for each cell recorded on the probe. The broad RFs identified in all 33 recording sessions from the two monkeys are depicted on the SC map (Fig-1E), which shows that we have recorded neurons situated in the middle and caudal portions of the colliculus.

### Behavioral Task

The monkeys were trained to perform visually guided saccades (Fig-1A) to single objects appearing in the RF of the neurons or in a position 180-degree opposite (AF) to the RF. These single-object trials were interleaved with two object choice trials that occurred in pseudo-random order within a block. A single block consisted of 128 trials with equal valued choices, unequal valued choices, and single-object trials in the RF and AF. The monkey performed 2-3 blocks of trials in a recording session. This paper focuses exclusively on the behavior and neuronal data obtained on single-object saccade trials presented in the RF of the neurons.

A single stimulus set is composed of eight fractal objects, with four fractals arbitrarily designated as good objects (associated with high reward volume) and four as bad objects (associated with lower reward). Juice volumes were controlled by keeping the solenoid valves open for 250ms (good objects) or for 50ms (bad objects). The task began with a fixation spot (green-colored square) appearing at the center of the screen. The monkeys maintained their gaze at the central fixation spot for 300ms, after which visual targets appeared at the RF or AF locations. The animals were trained to direct their gaze to the target and hold it there for 500 ms, at which point an audio cue and juice reward delivery occurred. Diluted juice was used as positive reinforcement. The trial was aborted if the animals broke fixation during the pre-target (300ms) or post-saccade (500ms) interval. The same conditions were repeated immediately following an aborted trial to ensure a comparable number of correct trials in each block.

The monkeys were trained on each fractal set for more than ten days prior to the neural recording. This consistent long-term exposure will result in the formation of long-term, stable value memory that boosts the animals’ performance. The monkeys correctly chose the good objects more than 98% of the times when good and bad objects were presented simultaneously, suggesting that they could readily associate value with a large number (8 objects *4 sets = 32) of objects.

## Data Analysis

### Preprocessing

The neural data acquired through the Plexon system was processed using the MATLAB-based package KILOSORT (Pachitariu et al., 2016) to perform automated spike sorting on the signals acquired from 24 independent V-probe channels. We manually curated the auto-detected units using PHY (https://github.com/cortex-lab/phy), a software that allows visualization of identified units, merging similar units into one, or splitting the activity of one identified unit into different units. A total of 854 units were identified and manually verified in this manner from 33 sessions recorded from two different animals.

### Behavioral analysis

We considered an object fixated when the eye position was stationary and within 8° of the center of the object based on horizontal and vertical eye position traces. Saccade onsets were defined as the time point when the instantaneous velocity crossed 30 degrees/s, and the end of the saccade was marked when the velocity fell below this threshold. Saccades interrupted by blinks were omitted by using a velocity threshold of 700 degrees/s. Different saccade parameters, such as amplitudes, peak velocity, reaction time (RT), etc., were computed for further analysis.

Targets were placed at different screen positions with amplitudes ranging from 3 degrees to 25 degrees on either the left or right visual fields based on the RFs of the neurons that were being recorded in each session. Hence, the saccades had to be normalized to compare the velocities and amplitudes across sessions, which was accomplished by normalizing saccade durations (Vasudevan et al., 2023). Normalization involved reestimating the average saccade displacement at every 5% of the saccade duration such that each session had 20 data points going from 0 to 100%, irrespective of the number of data points in the original duration. A polynomial interpolation was used to estimate the saccade displacements at each of the 20 data points, and the same function was differentiated to calculate the angular velocities at the same points. This normalization was carried out separately for high-valued good and low-valued bad object trials.

### Neural Analysis

Spikes of well-isolated neural units identified by the KILOSORT algorithm were used for all analyses. Spike density functions (SDF) were generated using Gaussian kernels with a bandwidth of 10 ms after aligning the activity on the target onset or saccade onset. Neurons were included in the analysis if they had at least five valid trials from each task condition and also fired more than 20 spikes in the analysis time window.

### Functional classification of SC neurons

The activity of individual SC neurons evoked by single objects in the RF was used to classify cells into functional categories. Windows corresponding to baseline activity (-200 to 0ms), <u>Phasic</u> activity #1 (0 to 80ms), and <u>Phasic</u> activity #2 (80 to 160 ms) were defined relative to target onset (Fig-1D left). Visual activation was defined as a significant increase in the phasic#1 and phasic#2 activity compared to baseline. Windows corresponding to pre-saccadic (-100 to-50ms) and peri-saccadic (-50ms to 20ms) intervals were defined relative to saccade onset (Fig-1D right). Motor activity was defined as a significant difference between pre-and peri-saccadic firing rates. A Kruskal-Wallis test was used to assess statistical significance between the firing rates in each window.

Visual neurons have a significantly higher activation during phasic#1 and phasic#2 compared to the baseline activity when aligned to the target onset. In addition, when aligned on saccades, there is no significant difference between pre-and peri-saccadic activity. Cells that fulfilled these criteria were defined as visual neurons.

Motor neurons are cells that show a significant difference between activities in the pre-saccadic and peri-saccadic intervals. In addition, when aligned on target onset, the phasic#1 activity is significantly lower than the phasic#2 activity. Cells that fulfilled both these conditions were defined as motor cells.

Visuomotor cells have a significant activation during phasic#1 and phasic#2 compared to the baseline activity when aligned on target onset. In addition, when aligned on saccades, there is a significant difference between the pre-and peri-saccadic intervals. Cells that fulfilled both these criteria were defined as visuomotor neurons.

Tonic neurons are defined as cells that showed significant activity reduction during the phasic#1 or phasic#2 compared to the baseline. In contrast to visual and motor cells, tonic neurons have high baseline firing rates that are suppressed by target onset in the RF.

520 units out of the total 854 identified units were classified into four functional subtypes using the above criteria. The rest 334 units were not classified and excluded from subsequent analysis because 1) the cells did not respond to the task in the manner specified above, 2) the cells responded in less than 5 trials, or did not fire more than 20 spikes during the task (n=170) 3) the cells were task modulated but did not meet the statistical criteria used for classification in this study (n=164).

### Reward Modulation

We studied the effect of reward in each functional subtype of SC neurons separately. To visualize this modulation, SDFs were generated separately for trials when good and bad objects were presented in the neurons’ RF. To quantify neuronal modulation by object value, we measured each neuron’s response to 4 different good objects by summing the number of spikes within the window 80-240ms after object presentation. We then compared it to the response of the same neuron in the same window when four bad objects were presented in the RF of the neuron. The statistical significance of this neuronal modulation was assessed using the Wilcoxon signed-rank test. Additionally, AUROC values were computed using the neural activity in the time interval 80-160ms after target onset to parametrize the reward modulation into a uniform scale ranging from 0-1. Only neurons that showed significant AUROC values were used in the analysis shown in Fig-9.

We also determined the time course of value modulation. To determine when the differences between good and bad objects became significant, we initially computed the difference between the firing activities of these two conditions (SDF_good_-SDF_bad_). We then computed the mean and the variability of this difference during the baseline period (-300 to 0ms). These measures were used to identify a threshold (mean + 3 SD) for each neuron separately. The time when the difference between activity profiles of good and bad objects crossed this threshold and stayed above the threshold continuously for 20ms was identified. This time point was considered as the onset of the value of modulation of that particular neuron. Using this approach, the onset of value modulation was determined for each cell separately, and the mean onset times were compared between the different subtypes.

We also applied the same onset detection procedure on the averaged population responses of each neuronal subtype. Compared to the mean onset times determined from individual neurons, the onsets detected from the population responses were approximately 10 ms earlier.

### Statistical tests

For population analysis, firing rates, quantified separately for each neuron, in each condition is pooled together, and a paired Wilcoxon signed-rank test was used for statistical comparisons between good and bad object responses. Error bars in all plots show the standard error of the mean (SEM) unless otherwise noted. Significance thresholds for all tests in this study were α=0.05. The means and the p-values are tabulated in the supplementary tables.

### Bootstrapping Procedure

The experiment performed in this study allowed us to directly assess the effect of reward on the response of different types of SC neurons. It is unclear if this modulation affected the visual or the pre-saccadic motor processes encoded in the SC neurons. To test this systematically, we employed a bootstrapping procedure (Supplimentry Fig-2) on the data. The bootstrapping procedure is based on the idea that a single neuron does not control behavior in individual trials, but rather, a population of neurons pools their activity to control behavior. For example, in the above experiment, multiple visual neurons respond to an object in a particular location on the visual field and contribute to the visual encoding of targets. Similarly, many motor and visuomotor neurons pool their activity to direct the saccade into the object’s location. The following steps were used to bootstrap the experimental data separately for each functional subtype of SC neurons.

1) The trials in each experimental session were divided into three quantiles (SupFig-2A) based on the saccadic metrics of reaction time (RT). Neuronal activities recorded for each neuron were likewise divided into three quantiles (SupFig-2B) based on the same behavioral quantiles.
2) To simulate the neural activity for a hypothetical trial belonging to one of the quantiles, the neuronal activity corresponding to individual trials belonging to the same behavioral quantile is pooled together from 25 random neurons. Based on this pooled activity, SDF corresponding to the hypothetical trial was generated (SupFig-2C).
3) 10-35 trials were simulated separately (SupFig-2D) for each behavioral quantile, and the average activity across trials was computed. The average firing rates during multiple epochs is simulated in this manner for each quantile (SupFig-2E) and compared to identify the effect of behavioral parameters on SC neurons. These steps constitute a single repeat of the bootstrapping process

This bootstrapping procedure was repeated 10000 times, and the mean firing rate/time to peak activity and its 95% confidence intervals are identified and tabulated in the supplementary tables. This analysis was also performed separately on good and bad object trials to test the effect of reward.

### Correlation Analysis

We computed correlation coefficients to assess the linear relationship between saccade behavioral parameters and the firing activity of well-isolated SC neurons. The observed neural activity was aligned on target onset, and the average firing rate in the time window of 80 to 160 ms was calculated for each trial. Only trials with at least 5 spikes were included in this analysis. The Pearson correlation coefficient was computed between the average firing rate in each trial and the corresponding behavioral measure (RT or PV). Only those neurons with at least 15 valid trials are included in this analysis. The calculated correlation was used as an index to quantify the sensitivity of SC neurons to the upcoming saccade behavior. We computed two metrics for each identified neuron: the Reaction Time Index (firing rate vs. RT correlation) and the Peak Velocity Index (firing rate vs. PV correlation). The correlation was considered significant if the p-value was less than or equal to 0.05.

## Supplementary Information: Titles and Legends

**Supplementary Figure 1: Value modulation in representative SC neurons**

Example neurons recorded from SC. Raster plots (top) and spike density functions (bottom) aligned on target onset (left) and saccade onset (right) are shown. Red and blue plots showed trials when a saccade was directed to good vs. bad objects, respectively. Example neurons were selected to illustrate the different functional subtypes of SC neurons, namely: **(A, B)** Visual neuron, **(C, D)** Visuomotor neuron, **(E, F)** Motor neuron, and **(G, H)** Tonic neuron.

**Supplementary Figure-2: Schematic of the bootstrapping procedure.**

**(A)** The distribution of reaction time observed during the saccade task in each session is shown with trials grouped into three quantiles. **(B)** Firing rate histograms and raster plots of all recorded neurons are likewise divided into three groups according to behavioral quantile. **(C)** Simulated single-trial responses were generated by pooling 25 randomly selected responses from all recorded neurons. **(D)** 10-35 trials are simulated for each behavioral quantile. Population activity is generated by averaging over these simulated trials drawn from each behavioral quantile. **(E)** This bootstrapping is repeated 10,000 times to generate the averaged activity profiles, which show a clear effect of saccade RT on SC neurons.

**Supplementary Figure-3: Correlation between firing rate and saccade behvaior.**

A scatter plot between neuronal activity in the time window 80-160 and the saccade RT in the same trial is shown separately for a representative visual **(A)**, visuomotor **(B)**, motor **(C),** and tonic

**(D)** neuron. The dashed line shows the best-fit line. The number on the top of each panel indicates Pearson’s correlation coefficient and the associated p-value. A scatter plot between neuronal activity in the time window 80-160 and the saccade PV in the same trial is shown separately for the same representative visual **(E)**, visuomotor **(F)**, motor **(G),** and tonic **(H)** neurons. The rest of the conventions are the same as above. The correlation was calculated in three separate epochs:

-100 to 0 ms, 0 to 80 ms, and 80-160 ms while the activity was aligned on target onset. The number of neurons that showed a significant correlation to RT (black) and PV(grey) in each time epoch is shown as separate bars for the population of visual **(I)**, visuomotor **(J)**, motor **(K)**, and tonic **(L)** neurons.

**Supplementary Figure-4: Time course of value modulation in SC population activity.**

The time course of value modulation of SC neurons is represented as the average difference between the good and bad object activities aligned on target onset (SDF_good_-SDF_bad_) and separately calculated for Visual **(A)**, Visuomotor **(B)**, and Motor **(C)** neurons. The vertical dashed line indicates when the population value difference became significantly different from the baseline. The histogram of onset times estimated from individual cell responses is shown separately for Visual **(D)**, Visuomotor **(E)**, and Motor **(F)** neuron subtypes.

**Supplementary Table 1: Quantification of mean peak time in different phases of SC response.**

A table showing the mean peak time for the simulated population activity for each reaction time quantile for Visual, Visuomotor, and Motor neurons. The mean peak time in each phase (E_VIS_, L_VIS_, and PRE_SAC_) is separately tabulated. The values tabulated here are related to the analysis shown in Figure 5.

**Supplementary Table 2: Quantification of value modulation in different phases of SC response.**

A table showing the comparison of average firing activity between good and bad objects during the early visual (E_VIS_), late visual (L_VIS_), and pre-saccadic (PRE_SAC_) phases for Visual, Visuomotor, and Motor neurons. The firing rate comparison is done separately and tabulated in the three reaction time quantiles (short, medium, and long). The values tabulated here are related to the analysis shown in Figure 6.

