## Supplemental Information for "Multiple groups of neurons in the superior colliculus convert value signals into saccadic vigor"

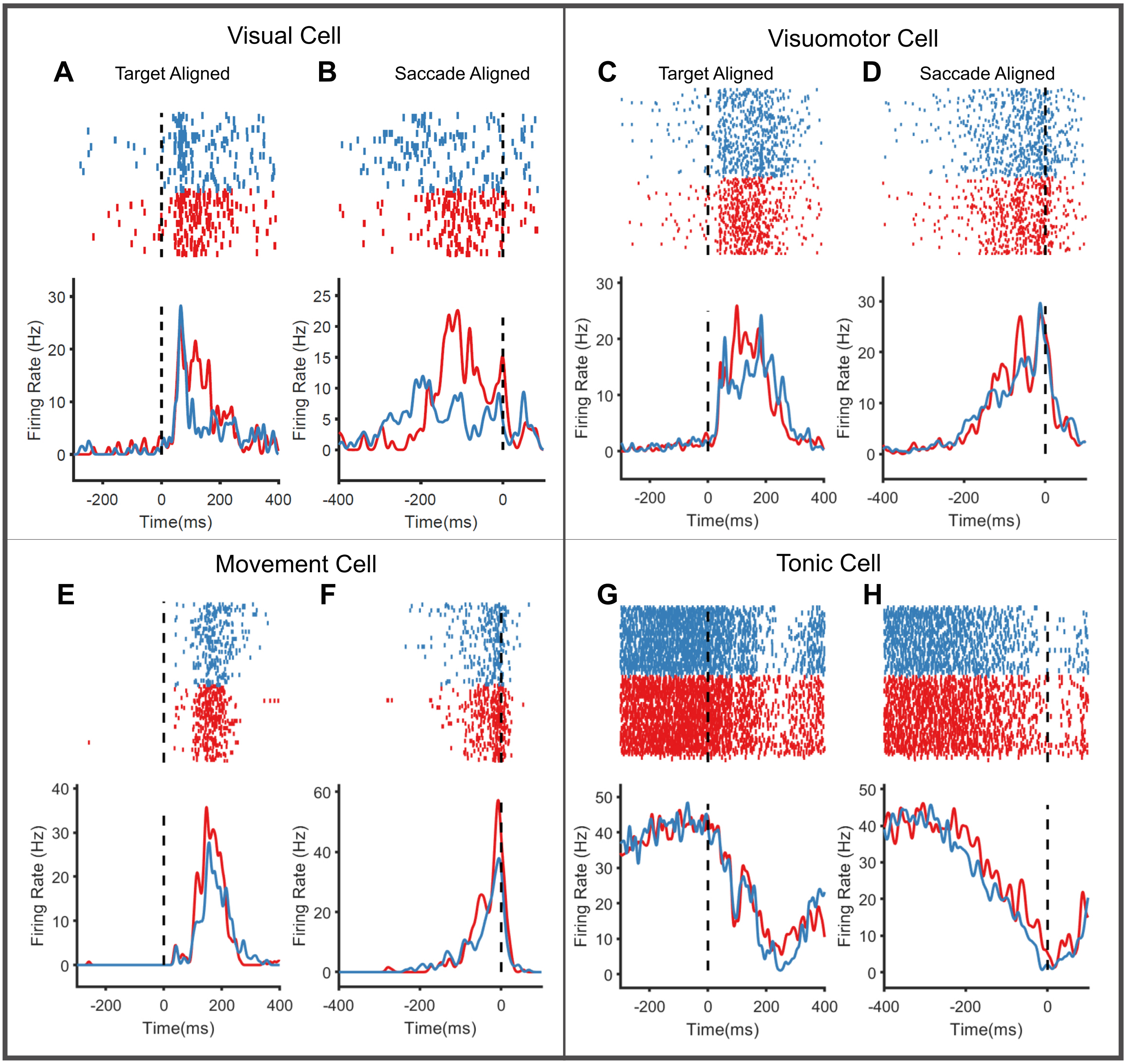

**Supplementary Figure - 1**

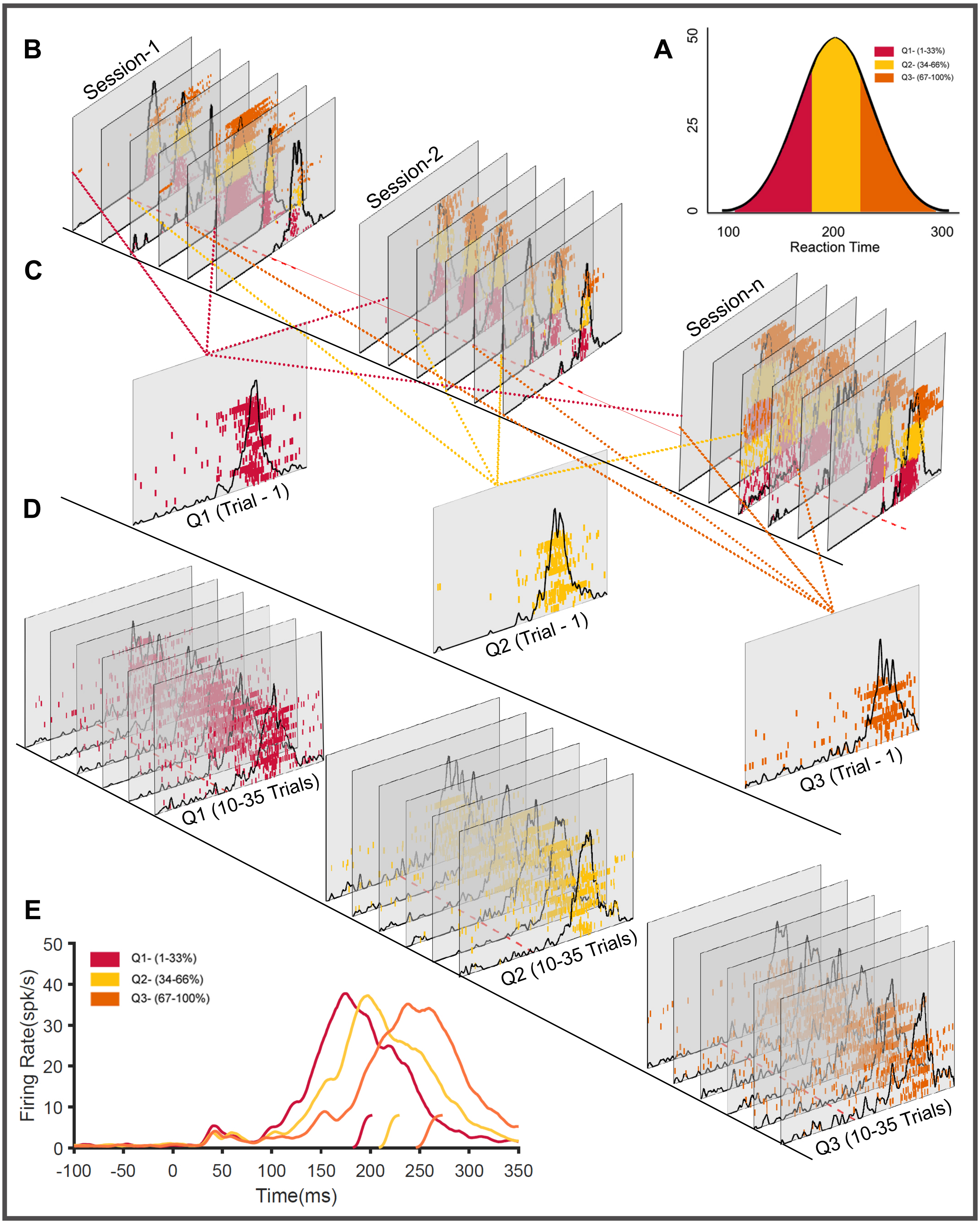

**Supplementary Figure- 2**

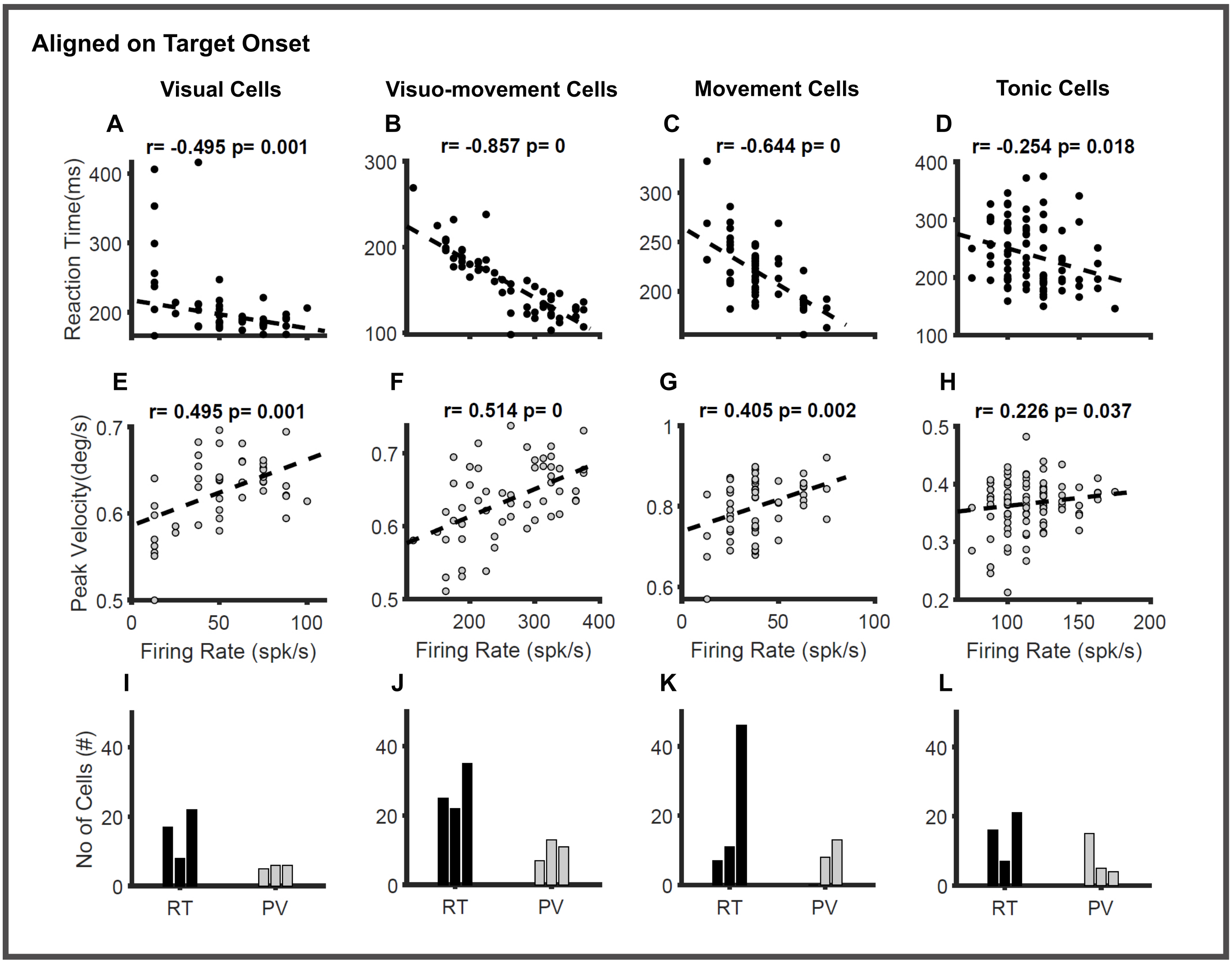

**Supplementary Figure - 3**

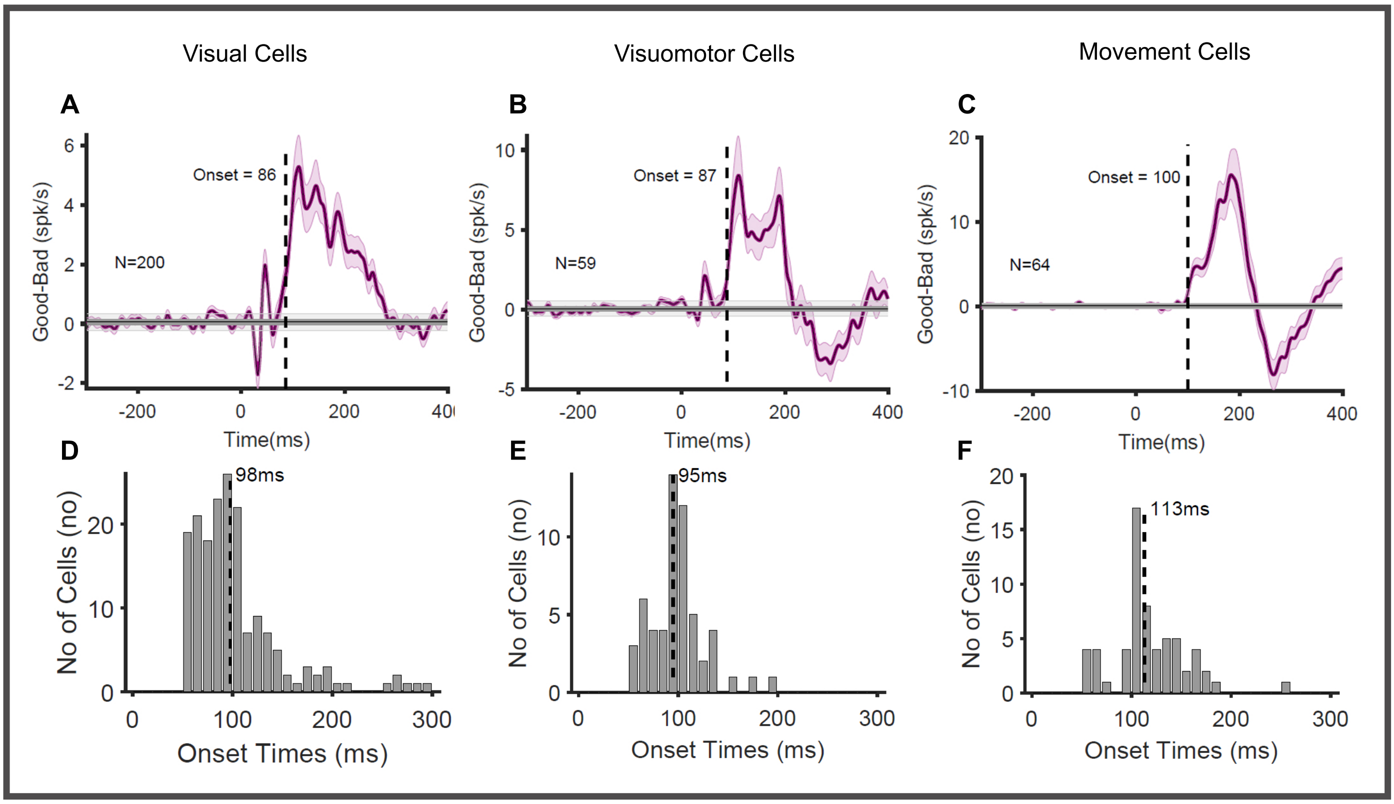

**Supplementary Figure - 4**

**Table-1**

| Neuron Subtype | Epoch | Time to Peak (ms) | | | | | | | | |
|---|---|---|---|---|---|---|---|---|---|---|
| | | Short | | | Medium | | | Long | | |
| | | **Mean** | 95% Intervals | | **Mean** | 95% Intervals | | **Mean** | 95% Intervals | |
| | $E_{VIS}$ | **42** | 39 | 45 | **43** | 40 | 47 | **43** | 40 | 47 |
| **Visual Neuron** | $L_{VIS}$ | **101** | 97 | 105 | **101** | 98 | 105 | **100** | 96 | 105 |
| | $Pre_{SAC}$ (Target) | **164** | 154 | 175 | **166** | 155 | 177 | **184** | 167 | 202 |
| | $Pre_{SAC}$ (Saccade) | **-86** | -100 | -75 | **-102** | -115 | -87 | **-130** | -148 | -109 |
| | $E_{VIS}$ | **39** | 38 | 41 | **40** | 39 | 42 | **40** | 39 | 41 |
| **Visuomotor Neuron** | $L_{VIS}$ | **109** | 105 | 112 | **110** | 105 | 114 | **109** | 106 | 113 |
| | $Pre_{SAC}$ (Target) | **154** | 147 | 164 | **165** | 157 | 174 | **193** | 183 | 201 |
| | $Pre_{SAC}$ (Saccade) | **-14** | -15 | -11 | **-13** | -16 | -10 | **-11** | -14 | -8 |
| | $E_{VIS}$ | **44** | 41 | 48 | **45** | 41 | 49 | **44** | 41 | 48 |
| | $L_{VIS}$ | **110** | 102 | 118 | **108** | 102 | 113 | **110** | 105 | 114 |
| **Motor Neuron** | $Pre_{SAC}$ (Target) | **165** | 158 | 171 | **186** | 180 | 191 | **216** | 207 | 225 |
| | $Pre_{SAC}$ (Saccade) | **-9** | -11 | -8 | **-8** | -9 | -7 | **-9** | -10 | -8 |

**Supplementary Table-1: Quantification of mean peak time in different phases of SC response.**

A table showing the mean peak time for the simulated population activity for each reaction time quantile for visual, visuomotor, and motor neurons. The mean peak time and the 95% confidence intervals obtained by bootstrapping in each phase ($E_{VIS}$, $L_{VIS}$, and $Pre_{SAC}$) are separately tabulated. The values tabulated here are related to the analysis shown in Fig-5

**Table- 2**

| Firing Rates (spk/s) | | | | | | | | | | | | | |
|---|---|---|---|---|---|---|---|---|---|---|---|---|---|
| **Subtype** | **Epoch** | **Short** | | | | **Medium** | | | | **Long** | | | |
| | | **Mean** | | **95% Interval** | | **Mean** | | **95% Interval** | | **Mean** | | **95% Interval** | |
| | | Good | Bad | Good | Bad | Good | Bad | Good | Bad | Good | Bad | Good | Bad |
| **Visual Neuron** | $E_{VIS}$ | 31.37 | 28.51 | 29.6-33.3 | 26.7-30.4 | 30.32 | 28.24 | 28.6-32.2 | 26.5-30.1 | 27.29 | 26.51 | 25.5-29.1 | 24.8-28.3 |
| | $L_{VIS}$ | 37.88 | 30.46 | 35.1-40.8 | 28.1-32.8 | 39.04 | 29.95 | 36.1-42.1 | 26.8-31.2 | 37.19 | 27.71 | 34.3-40.2 | 25.7-29.9 |
| | $Pre_{SAC}$ | 29.18 | 23.65 | 26.6-31.8 | 21.5-25.9 | 30.76 | 23.08 | 28.3-33.4 | 21.2-25.1 | 30.00 | 20.07 | 27.5-32.7 | 18.33-21.9 |
| **Visuomotor Neuron** | $E_{VIS}$ | 30.23 | 27.64 | 28.0-32.6 | 25.6-29.8 | 27.53 | 26.58 | 25.6-29.6 | 24.7-28.5 | 25.55 | 27.03 | 23.6-27.5 | 25.0-29.1 |
| | $L_{VIS}$ | 51.21 | 43.62 | 47.5-55.0 | 40.4-46.9 | 46.02 | 35.10 | 42.7-49.4 | 32.3-37.9 | 38.60 | 33.66 | 35.6-41.7 | 31.0-36.3 |
| | $Pre_{SAC}$ | 57.76 | 48.45 | 53.9-61.7 | 44.6-52.3 | 54.63 | 42.22 | 50.8-58.6 | 39.1-45.3 | 46.35 | 38.56 | 43.1-49.8 | 35.7-41.5 |
| **Motor Neuron** | $E_{VIS}$ | 10.42 | 10.68 | 9.2-11.7 | 9.4-12.0 | 8.83 | 8.43 | 7.7-10.1 | 7.2-9.7 | 8.64 | 10.01 | 7.4-10.0 | 8.6-11.6 |
| | $L_{VIS}$ | 34.86 | 29.40 | 32.1-37.7 | 26.8-32.0 | 27.26 | 21.50 | 24.8-29.8 | 19.4-23.7 | 20.60 | 19.49 | 18.6-22.7 | 17.5-21.6 |
| | $Pre_{SAC}$ | 78.33 | 64.47 | 73.0-83.9 | 59.6-69.5 | 71.81 | 50.34 | 66.3-77.4 | 46.5-54.5 | 53.83 | 36.85 | 49.5-58.5 | 33.7-40.1 |

**Supplementary Table-2: Quantification of value modulation in different phases of SC response.**

A table showing the average firing activity of good and bad object trials during the $E_{VIS}$ (40-80 ms), $L_{VIS}$ (81-130ms), and $Pre_{SAC}$ (131-170ms) phases of visual, visuomotor, and motor neurons. The mean firing rate and the 95% confidence interval obtained from the repeated bootstraps are tabulated separately for the three reaction time quantiles (short, medium, and long). The values tabulated here are related to the analysis shown in Fig-6
